# Structural basis for L-isoaspartyl-containing protein recognition by the PCMTD1 cullin-RING E3 ubiquitin ligase

**DOI:** 10.1101/2025.05.21.654933

**Authors:** Eric Z. Pang, Boyu Zhao, Cameron Flowers, Elizabeth Oroudjeva, Jasmine B. Winter, Vijaya Pandey, Michael R. Sawaya, James Wohlschlegel, Joseph A. Loo, Jose A. Rodriguez, Steven G. Clarke

**Affiliations:** Department of Chemistry and Biochemistry, University of California, Los Angeles, Los Angeles, CA 90095, USA; Department of Biological Chemistry, University of California, Los Angeles, Los Angeles, CA 90095, USA; UCLA-DOE Institute, University of California, Los Angeles, Los Angeles, CA 90095, USA; Molecular Biology Institute, University of California, Los Angeles, CA 90095, USA

**Keywords:** Aging, L-isoaspartyl, E3 ligase, cullin-RING ligase, CRL, Methyltransferase, *S*-adenosylmethionine, Native mass spectrometry, Cryo-EM

## Abstract

A major type of spontaneous protein damage that accumulates with age is the formation of kinked polypeptide chains with L-isoaspartyl residues. Mitigating this damage is necessary for maintaining proteome stability and prolonging organismal survival. While repair through methylation by PCMT1 has been previously shown to suppress L-isoaspartyl accumulation, we provide an additional mechanism for L-isoaspartyl maintenance through PCMTD1, a cullin-RING ligase (CRL). We combined cryo-EM, native mass spectrometry, and biochemical assays to provide insight on how the assembly and architecture of PCMTD1 in the context of a CRL complex fulfils this alternative mechanism. We show that the PCMTD1 CRL complex specifically binds L-isoaspartyl residues when bound to AdoMet. This work provides evidence for a growing class of E3 ubiquitin ligases that recognize spontaneous covalent modifications as potential substrates for ubiquitylation and subsequent proteasomal degradation.

**eTOC Blurb:** Limiting the accrual of L-isoaspartyl damaged proteins is essential during aging. While this is thought to be mediated solely by the repair activity of the protein, PCMT1, Pang *et* al. now demonstrate that a related protein, PCMTD1, functions as a cullin-RING ligase to selectively target L-isoaspartyl-damaged substrates for potential regulation by the ubiquitylation-proteosomal system.

**Highlights:**

- Atomic cryo-EM structure of CRL5-PCMTD1 determined
- Architecture of PCMTD1 when complexed as a CRL supports ubiquitylation activity
- PCMTD1 recognizes L-isoaspartyl residues as a recruitment motif for potential CRL activity
- Recognition of L-isoaspartyl residues is dependent on cofactor engagement

## Introduction

Molecular aging is marked by the accumulation of covalently modified polypeptides^1^. Survival during aging therefore depends in part on limiting the accumulation of proteins damaged by spontaneous chemical reactions to maintain homeostasis^1,2^. Examples of such damages include the non-enzymatic modification by the oxidation of various residues^3^, the glycation of lysine residues^4^, and the deamidation and isomerization of asparagine and aspartyl residues into L-isoaspartyl residues^5^. To date, enzymatic recognition of these damages followed by subsequent repair has been the most well-characterized mechanism for regulating age-related damages which include the action of methionine sulfoxide reductases^6^, ketoamine repair enzymes^7^, and the protein-L-isoaspartyl (D-aspartyl) O-methyltransferase (PCMT1)^8^.

Protein homeostasis is also maintained by the equilibrium of protein synthesis and protein turnover to maintain a functional proteome; the majority of that protein turnover is regulated by the ubiquitin-proteosome system (UPS)^9,10^. Within the UPS, E3 Ubiquitin (Ub) ligases facilitate the degradation of proteins by recognizing specific targets. This recognition is mediated by substrate-specific sequence motifs known as degrons^9–11^. Degron binding can be further regulated by enzyme-catalyzed post-translational modifications (PTMs) that either promote or inhibit substrate∼E3 Ub ligase interactions^12^. However, evidence is sparse for a general class of E3 Ub ligases that recognizes degrons defined by non-enzymatically-derived PTMs^13^.

Of the E3 Ub ligases, cullin-RING ligases (CRLs) represent the largest subclass of such enzymes, and their ubiquitylation activity is also influenced by degron PTMs. CRLs feature a conserved catalytic core formed by a cullin protein and a RING protein^14^. Acting as scaffolds, cullins link the RING protein at its C-terminus with a diverse pool of substrate receptors at its N-terminus^14^. Receptors are able to bind specific substrates while interacting with cullins through specific motifs^15^. For CUL5 CRLs, these substrate receptors consist of suppressor of cytokine signaling (SOCS)-box proteins that utilize an Elongins B/C heterodimer (ELOBC) to associate with the N-terminus of CUL5 via the BC-box and Cul-box motifs, respectively^16,17^. Ubiquitylation of targets is driven by the spatial and temporal coordination of receptor-bound substrates with ubiquitylation machinery recruited by RING proteins^15^.

Similar to other E3 Ub ligases, ubiquitylation activity by CRLs can be further tuned through covalent PTMs of CRL degrons which can increase substrate diversity and enhance substrate specificity. While enzyme-catalyzed PTMs such as phosphorylation are viewed as the primary sources of covalent modifications to CRL degrons, those defined by non-enzymatic PTMs have also been suggested through recent studies of cereblon (CRBN)^18,19^ and the PCMTD-proteins^16^, which may position CRLs in age-related proteomic maintenance. Notably, the L-isoaspartyl damage requires the initial generation of an unstable succinimide intermediate, a five-membered cyclic imide, which is then non-enzymatically hydrolyzed and completely depleted to either L-aspartyl, or to a greater extent, the L-isoaspartyl damage^20^.

PCMTD1 and PCMTD2 have been proposed to recognize spontaneously isomerized and racemized aspartyl and asparagine residues, including the L-isoaspartyl modification, formed in processes that do not induce protein backbone cleavage^16^. Because PCMTD1 retains an N-terminal PCMT-like domain, it binds the methyl donor cofactor, S-adenosylmethionine (AdoMet), but does not methylate L-isoaspartyl-containing substrates^16^. Through its C-terminal domains, PCMTD1 has been shown to interact with the CUL5-RBX2 cullin catalytic core through ELOBC to form a putative CRL complex, CRL5-PCMTD1, *in cells* and *in vitro*^16^. Given these results, we proposed PCMTD1 may mediate L-isoaspartyl damages not through enzymatic repair, but through substrate ubiquitylation and possible proteasomal degradation.

Using single particle cryo-EM, we now reveal the architecture of CRL5-PCMTD1, which parallels that of active CRLs. This structure reveals key structural motifs that facilitate complex assembly and the specific architecture of PCMTD1, including features that abolish its methyltransferase activity. We also demonstrate through native mass spectrometry (nMS) and biochemical assays that L-isoaspartyl damage can act as a recognition site for substrate engagement by PCMTD1, which unexpectedly requires co-engagement of AdoMet for substrate engagement. Overall, these results suggest PCMTD1 belongs to a novel class of E3 Ub ligases responsible for regulating proteins damaged by non-enzymatic covalent damages and functions complementary with PCMT1 in mitigating the accrual of L-isoaspartyl-damaged proteins.

## Results

### PCMTD1 assembles a cullin-RING ligase complex

Recent studies have suggested that PCMTD1 can interact with CUL5-RBX2 through ELOBC to form a CRL5-PCMTD1 complex^16^. To clarify the overall composition, stoichiometry and arrangement of subunits in this complex *in vitro*, we co-expressed and purified PCMTD1 and ELOBC, and mixed this complex with recombinantly expressed CUL5 and RBX2, forming the CRL5-PCMTD1 complex (Fig. 1A, Sup. Fig 1A). We first assessed the structural integrity of the sample by gel filtration and nMS^21,22^, which indicated the complex was pentameric, containing one subunit of each protein component from PCMTD1-ELOBC and CUL5-RBX2 (Fig. 1B). This was supported by negative stain electron microscopy that revealed a horseshoe shaped particle that accommodated an AlphaFold model of PCMTD1-ELOBC (Sup. Fig 1B) and indicated the complex might be amenable to single particle cryo-EM characterization.

**Figure 1.**
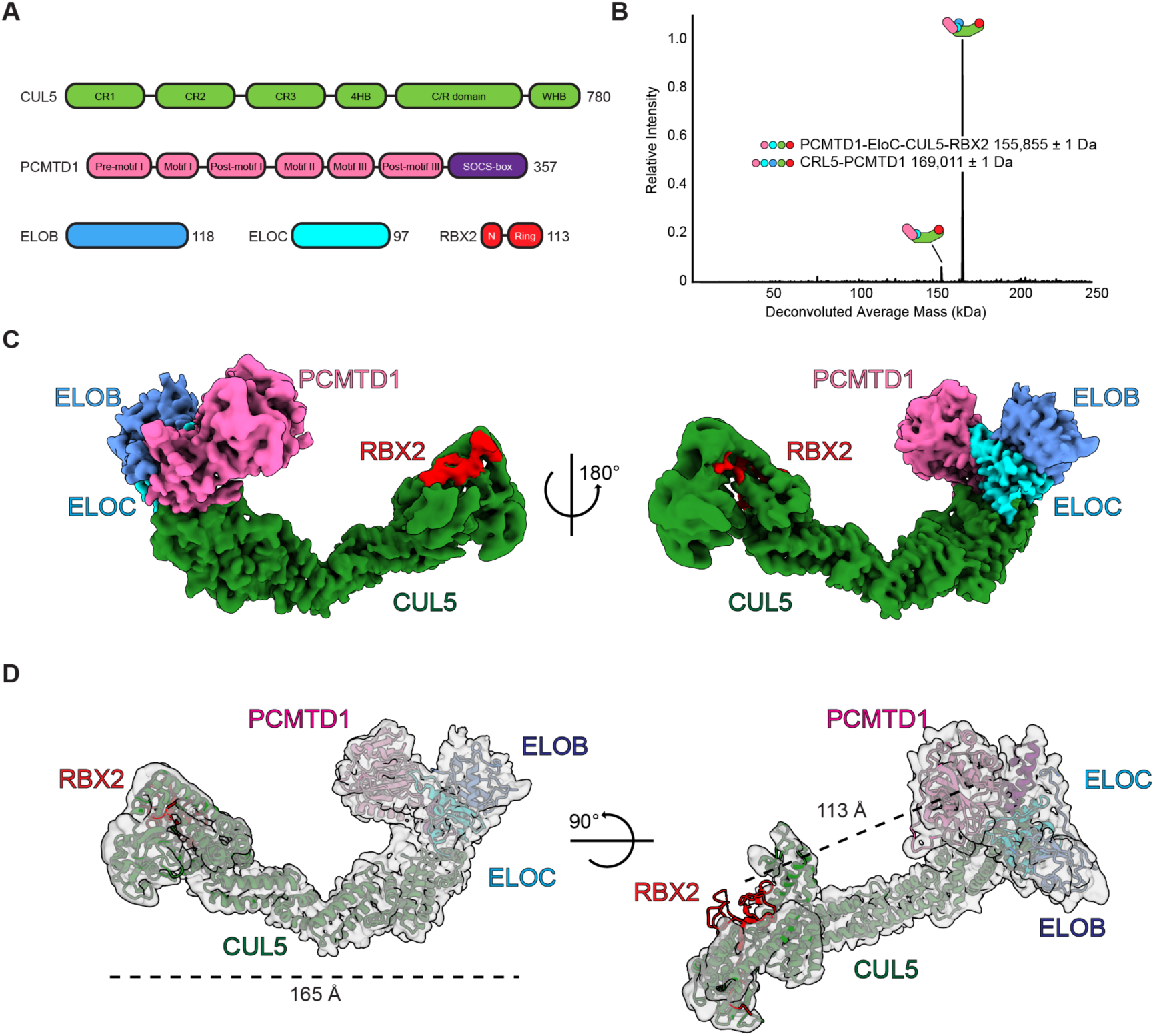
The structure of CRL5-PCMTD1. A) Schematic representation for all protein subunits used to form the CRL5-PCMTD1 complex. B) Native (deconvolved) mass spectrum of CRL5-PCMTD1 after *in vitro* reconstitution. C) Two orientations of a composite map of CRL5-PCMTD1 reconstructed from cryo-EM. Electron density for each subunit has been colored according to the schematic in panel A. D) The determined model of CRL5-PCMTD1 within the cryo-EM map used for structure determination with the span of the complex (left) and distance from PCMTD1 and RBX2 (right) marked. See Sup. Fig. 1 for sample preparation details of CRL5-PCMTD1. See Sup. Fig. 2 for cryo-EM structural characterization details. See also Sup. Fig. 3 for additional native and native top-down mass spectrometry data of CRL5-PCMTD1.

Single particle cryo-EM of CRLs can be challenging due to their aspect ratio making them prone to preferred-orientation, complex dissociation, and air-water interface damage^23–27^. To mitigate these challenges, we used the Chameleon and its self-wicking grids to control relative ice thicknesses during sample vitrification to mitigate these effects and to avoid the need to cross-link the multi-subunit CRL5-PCMTD1 complex (Sup. Fig. 1B)^28,29^. With these strategies, we obtained a 4.14 Å map of this complex that allowed structural determination of CRL5-PCMTD1 (Fig. 1C). Because the structure of PCMTD1 was previously unknown, we leveraged nMS to ensure the configuration of subunits in the CRL5-PCMTD1 complex. After storage at -80 °C, we found that the complex dissociates to subcomplexes, which are PCMTD1 bound to ELOB and ELOC while its interaction with RBX2 appears indirect because we do not observe PCMTD1-RBX2 subcomplexes (Sup. Fig 3A). This allowed us to unambiguously dock a predicted model of PCMTD1 into our cryo-EM map as an initial template for model building following rigid-body docking of previously determined structures of the other components within CRL5-PCMTD1 (Fig. 1D)^23,30–33^. This combination of nMS coupled with cryo-EM allowed us to reveal the positioning of each protein component in the CRL5-PCMTD1 complex, the interaction interfaces that maintain the complex, and key structural motifs in PCMTD1 responsible for cofactor engagement and potential substrate engagement.

CRL5-PCMTD1 adopts a horseshoe-like architecture, typical of CRLs^34–44^, that spans 165 Å in diameter and can be segmented into two-distinct regions, the substrate receptor complex (SRC) and cullin-scaffold (Fig. 1D, 2A)^45^. The SRC contains PCMTD1 bound to ELOBC (Fig. 2A) and is adjoined to the cullin-scaffold through the N-terminus of CUL5 at an interface stretching 23 Å while the C-terminus of CUL5 is bound to RBX2, harboring the RING domain needed for recruiting E2∼Ub conjugates (Fig. 2A,B)^46^. RBX2 appears flexible in the resolved map of CRL5-PCMTD1, with density evident only at a low map threshold, but appears to fully occupy this site based on nMS and native top-down MS (nTDMS) of the complex (Sup. Fig. 3, 4A)^47–49^. Despite its flexibility, RBX2 is positioned opposite and within the lateral plane of PCMTD1, primed to promote substrate ubiquitylation, as observed in other active CRLs (Sup. Fig. 4B)^38,39^.

**Figure 2.**
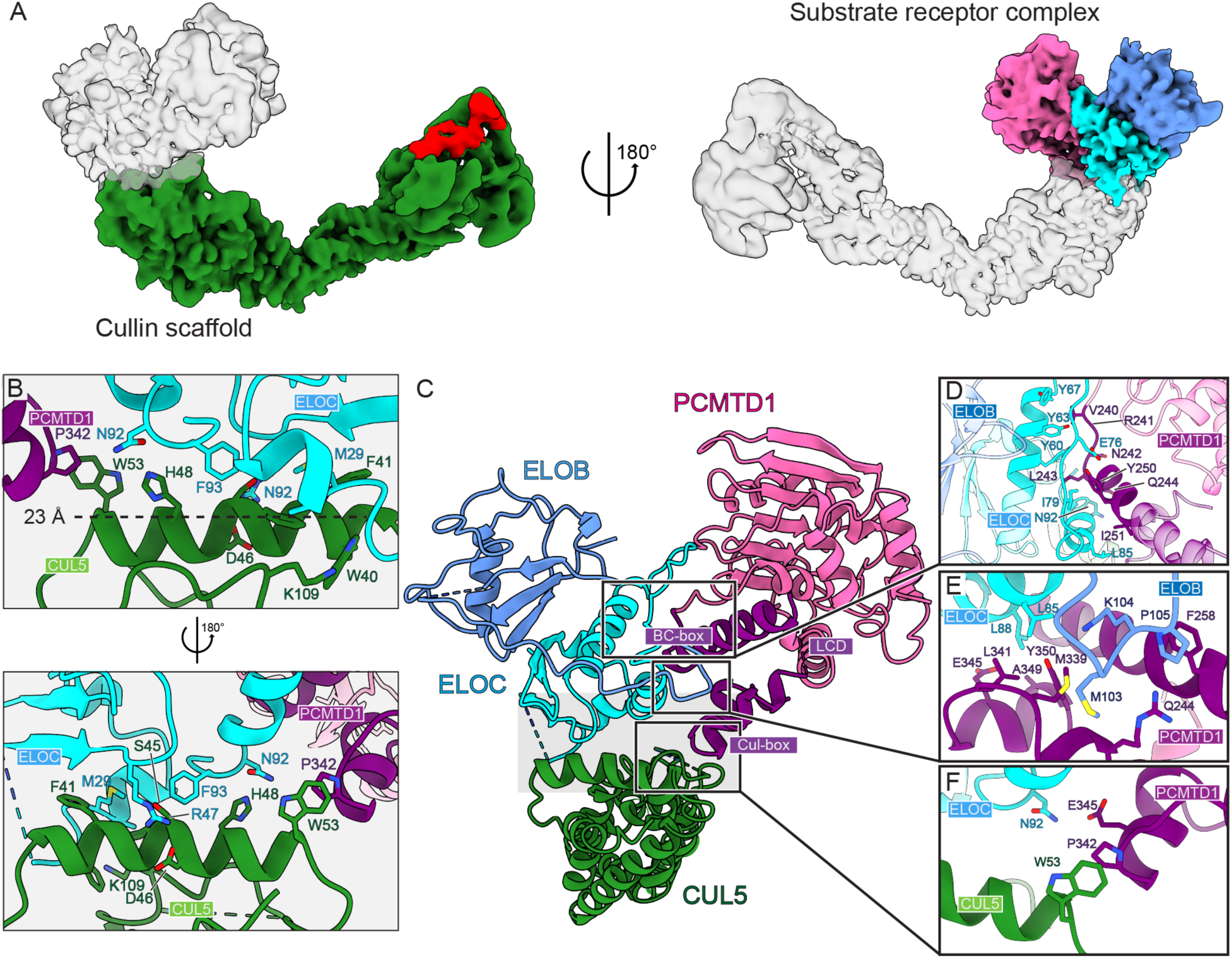
The interface between PCMTD1 and CRL components. A) Maps highlighting the cullin scaffold (left) and the PCMTD1 substrate receptor complex (right). B) Focused view of the SRC-cullin interface with the span of the interface marked. C) Ribbon model of the interfaces between PCMTD1 and its CRL partners. C) Focused view on interactions between the BC-box of PCMTD1 and ELOC. D) Interactions that promote the stabilization of the C-terminal tail of ELOB against PCMTD1’s ESB domain. E) The interface between the Cul-box of PCMTD1 and CUL5. A proximal H-bond interaction between ELOC and PCMTD1 is also shown. See also Sup. Fig. 4 for a comparison of the SRC interface to another ASB9, another SOCS-box protein, as well as a focused view of the PCMTD1 SRC-CUL5 interface from 3D reconstructions.

### PCMTD1 engages cullin-RING ligase components through both canonical and unique SOCS-box interactions

The BC-box and Cul-box motifs found in the C-terminal domain of PCMTD1 are essential for engaging ELOBC and CUL5^16,17,50^, but unlike from most SOCS-box proteins, these are separated by a longer 89 amino acid residue linker region (BC-Cul linker). This unusual architecture prompted close inspection of the PCMTD1 interfaces with CUL5, ELOB, and ELOC, compared to equivalent features in other CRL structures.

In line with other CUL5-binding SOCS-box proteins, PCMTD1 engages ELOC through its alpha-helical BC-box which is located within the C-terminal extended SOCS-box (ESB) domain of PCMTD1 (Fig. 2). Specifically, the carboxy terminus of R241 forms a hydrogen bond with Y60 of ELOC and residues V240, L243, Q244, Y250, and I251 found within the same face of an α-helix containing the BC-box of PCMTD1 form a hydrophobic interface against ELOC through residues Y60, Y63, Y67, E76, I79, and L85 (Fig. 2D).

PCMTD1 interacts with ELOC distinctly from the SOCS-box model, with an interface that extends beyond the BC-box on both its N-terminal and C-terminal sides and spans approximately 35 Å; this is ∼15 Å longer than the interface observed between ASB9 and ELOBC, a comparable SOCS-box SRC (Sup. Fig. 4C). ELOB further lengthens this interface by extending its C-terminal tail toward the PCMTD1 ESB domain (Fig. 2E). This large interface is uniquely facilitated by the first 14 residues of the BC-Cul linker in PCMTD1; extended α-helix geometry notably provides an additional platform for simultaneous engagement to ELOB and ELOC (Fig 2E). nMS confirms the presence of stable dimeric species of PCMTD1-EloB and PCMTD1-EloC, suggesting that Elongins B and C engage PCMTD1 in a robust and stable conformation (Sup. Fig. 3B, 3C).

In contrast to its interface with ELOBC, PCMTD1 interacts with CUL5 via a smaller interface, typical of SOCS-box proteins^23,30,51–54^, and facilitated by hydrophobic interactions between P342 of PCMTD1 and W53 of CUL5, and H-bond interactions between N92 of ELOC and E345 of PCMTD1 (Fig. 2F). Interestingly, our reconstruction reveals an unoccupied groove at the SRC-CUL5 interface measuring approximately 35 Å by 10 Å (Sup. Fig. 4D). This groove may provide the necessary clearance for dynamic processes associated with adapter protein exchange or substrate ubiquitylation, as previously reported in other CRL5 complexes^55^. This is supported by two key findings: (1) we do not observe stable PCMTD1-CUL5 heterodimers via nMS, and (2) nTDMS detects only N-terminal product ions of CUL5 following high-energy collisional dissociation (HCD) fragmentation of the complex (Sup. Fig. 3A, C)^48^.

### The N-terminal topology of PCMTD1 resembles the PCMT1 protein repair methyltransferase

Having characterized the PCMTD1 CRL complex, we next sought to determine the structural features underpinning its potential engagement of substrates for ubiquitylation. PCMTD1 consists of a two-domain structure containing an N-terminal 7-β-strand methyltransferase (7βS MTase) domain and its C-terminal ESB domain as mentioned above (Fig 3). The 7βS MTase domain of PCMTD1 comprises of a doubly wound α/β/α sandwich composed of six parallel β-strands (β1 – β5 and β7), an antiparallel strand (β6), and are surrounded by seven α-helices (αA, αB, αD, α1, α2, α5, and α7). This configuration is typical of enzymes in the protein-L-isoaspartate(D-aspartate) O-methyltransferase (PCMT) family^56^. The structure of PCMT1-class enzymes deviate only slightly from the general structure of AdoMet-dependent methyltransferases (where PCMT1 enzymes normally contain eleven α-helices), and this configuration allows specific binding and methylation of L-isoaspartyl residues for repair^8^.

**Figure 3.**
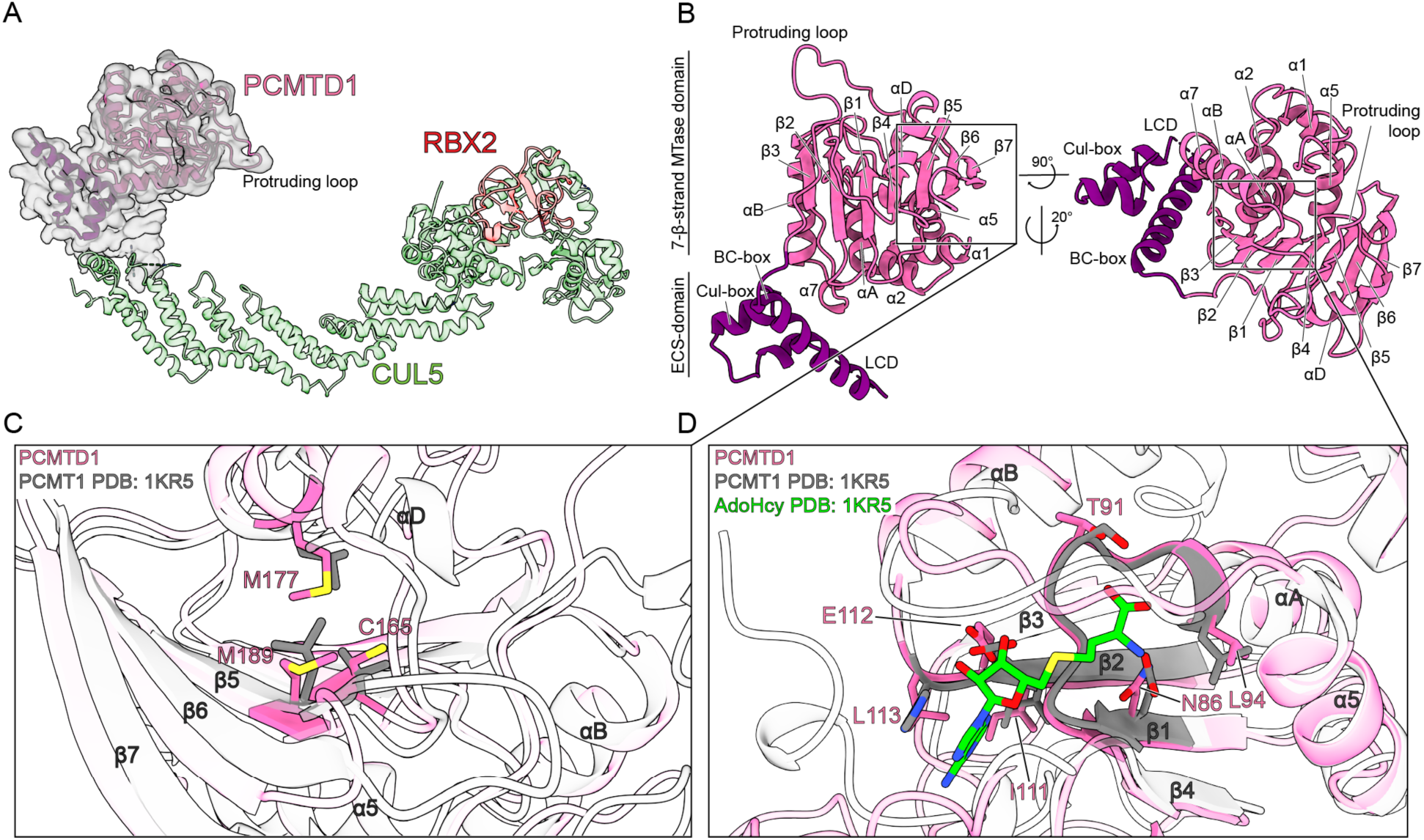
The structural topology of PCMTD1. A) The determined structure of PCMTD1 within its cryo-EM map highlighting its position within the CRL5 complex. B) Structure of PCMTD1 with secondary structure elements labeled. C) The novel Met-Cys-Met cluster in PCMTD1 with respective residues in PCMT1 superimposed. D) Critical residues in the cofactor binding site of PCMTD1 aligned with respective residues within the cofactor site of PCMT1. See Sup. Fig 5. for AdoMet binding and methylation assays of CRL5-PCMTD1.

In the context of PCMTD1-CUL5, PCMTD1 adopts a configuration that allows the putative cofactor and substrate binding sites found in its 7βS MTase domain to remain solvent accessible (Fig. 3). In contrast to PCMT, PCMTD1 harbors a novel protruding loop after β-strand 7 which is normally followed by a C-terminal α-helix (α-helix 9) in PCMT1. Although the function of α-helix 9 has not been characterized, the protruding loop feature of PCMTD1 interestingly points towards RBX2 in the CRL complex (Fig. 3B). PCMTD1 also contains within its twisted β-sheet a novel Met-Cys-Met cluster which is not found in PCMT1-class enzymes (Fig. 3C). The particular roles of the protruding loop and Met-Cys-Met features are unknown but may represent alternative uncharacterized functions PCMTD1 may harbor.

Even when complexed into a CRL, PCMTD1 still engages AdoMet (Sup. Fig. 5A), indicating that despite slight differences between the Rossman-like folds of PCMT1 and PCMTD1, the latter retains a functional Motif I and Post-motif I, responsible for AdoMet binding (Fig. 3D). Here, residues N86, L94, T91, E112, and L113, which are known to engage the adenosine and ribose moieties of AdoMet and AdoHcy through H-bonding and hydrophobic interactions, are oriented as in PCMT1 (Fig. 3D)^57^. However, while PCMTD1 engages AdoMet, it does not display methylation activity against isoaspartyl substrates^16^, nor does the CRL5-PCMTD1 complex (Sup. Fig. 5B).

### PCMTD1 specifically binds isoaspartyl-containing substrates in an AdoMet-dependent manner

Considering the similar structural topology between the PCMTD1 7βS MTase domain and PCMT1, the lack of methylation activity discussed above suggests that either PCMTD1 cannot engage L-isoaspartyl damaged proteins or engages substrates but lacks methyltransferase activity. To distinguish these possibilities, we evaluated substrate binding by PCMTD1-CUL5^tet^, a complex composed of PCMTD1 complexed with ELOB, ELOC, and CUL5^1–384^. We withheld RBX2 in this complex to avoid potential variability in nMS results caused by its metal binding properties^58^. nMS of this complex confirmed its native molecular weight, which we compared to that in the presence of peptides containing either L-aspartyl residues or L-isoaspartyl residues, reasoning that peptide recognition would reveal parental ions corresponding to PCMTD1-CUL5^tet^-bound peptides.

We employed two parent peptides of sequences KASADLAKY and VYPDLA, both well-characterized substrates of PCMT1^59^. We saw no evidence of PCMTD1-CUL5^tet^ binding to either L-aspartyl-containing peptide (Fig. 4C, Sup. Fig 6A). However, we found clear evidence for PCMTD1-CUL5^tet^ binding one molecule of each L-isoaspartyl containing peptide (Fig. 4C, Sup. Fig 6A). We only observed L-isoaspartyl peptide-dependent mass shifts when AdoMet was also present in the complex (Fig. 4C, Sup. Fig. 6A), suggesting that the AdoMet cofactor is required for isoaspartyl-peptide binding. Interestingly, we did not observe binding between the human PCMT1 repair methyltransferase and L-isoaspartyl containing peptides when performing similar nMS experiments (Fig. 4). Rather, the original L-isoaspartyl peptide was observed with a +14 Da mass shift, suggesting that upon AdoMet-dependent methylation, PCMT1 quickly disengages the methylated L-isoaspartyl product. In contrast, PCMTD1 stably engages L-isoaspartyl peptides in an AdoMet-dependent manner – the cofactor apparently acting to stabilize the complex with the isoaspartyl substrate rather than being a methyl donor source.

**Figure 4.**
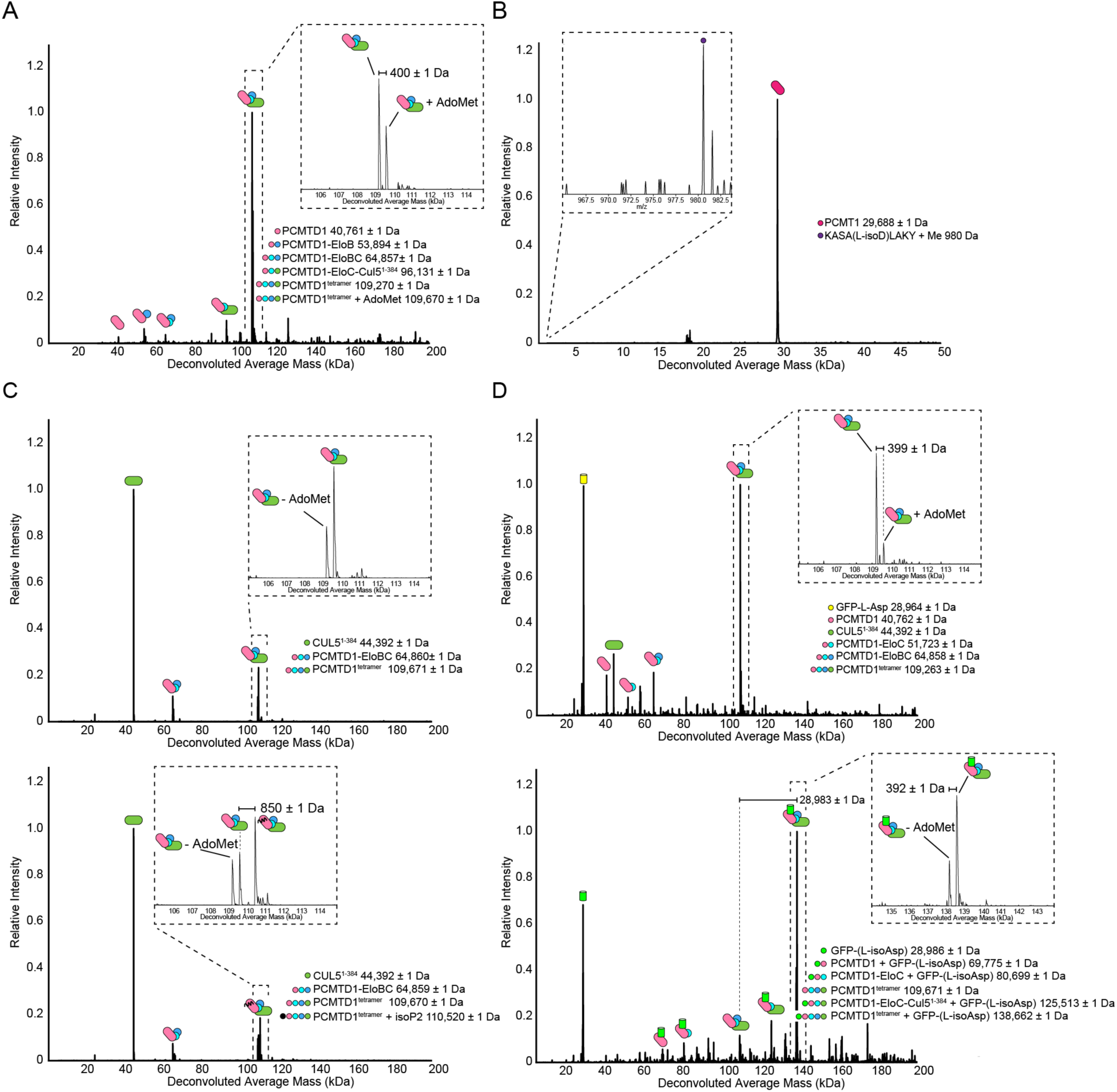
PCMTD1 binds L-isoaspartyl-containing peptides and proteins. A) Deconvolved native mass spectrum of PCMTD1-CUL5^tet^ (PCMTD1^tetramer^). B) Deconvolved native mass spectrum of PCMT1 spiked with AdoMet and peptide containing L-isoaspartyl. Inset represents raw spectra within the peptide’s m/z range. C) Deconvolved native mass spectra of PCMTD1-CUL5^tet^ with peptides containing L-aspartyl (top) and L-isoaspartyl residues (bottom). D) Deconvolved native mass spectra of PCMTD1-CUL5^tet^ spiked with sfGFP-Tet2 conjugated with peptides containing L-aspartyl (top) or L-isoaspartyl residues (bottom). Theoretical masses for all protein species are found in Table 2. See Sup. Fig. 6 for additional ligand binding with another peptide sequence and control experiments characterizing the generation of sfGFP-Tet2 peptide conjugates.

This led us to ask whether the PCMTD1 complex specifically recognizes L-isoaspartyl-containing proteins. By nMS, we determined whether PCMTD1-CUL5^tet^ could recognize a full-length GFP covalently linked to and presenting a solvent-exposed L-aspartyl or L-isoaspartyl-containing peptide^60^. We leveraged non-canonical amino acid incorporation of a reactive tetrazine amino acid into sfGFP for peptide conjugation with L-aspartyl or iso-aspartyl peptides harboring a reactive N-terminal trans-cyclooctene (sfGFP-L-Asp; sfGFP-L-isoAsp) and observed successful formation of these sfGFP neosubstrate by nMS (Sup. Fig. 6B). We found that PCMTD1-CUL5^tet^ is only able to bind sfGFP-L-isoAsp but not sfGFP-Tet2 or its clicked form to the non-isomerized peptide (Fig. 4D). Additionally, the relative amount of AdoMet-bound PCMTD1-CUL5^tet^ also increased as a result of binding sfGFP-L-isoAsp (Fig. 4D).

### PCMTD1 may lack methyltransferase activity due to the relative position of AdoMet and the isoaspartyl alpha-carboxyl group

The demonstrated binding of isoaspartyl-containing peptides and proteins to the PCMTD1 complex (Fig. 4C,D), the similarity of the PCMTD1 AdoMet binding pocket to that of human and *P. furiousus* PCMT1, and the biochemical evidence for AdoMet binding (Fig. 3D), suggest that the lack of methyltransferase activity of PCMTD1 arises from differences in the relative orientations of the alpha-carboxyl group of the isoaspartyl-peptide and the methyl group of AdoMet (Sup. Fig. 7)^56,61^. Methylation occurs only when the L-isoaspartyl residue aligns near the AdoMet sulfonium and this stringent alignment between substrate and cofactor is conserved broadly amongst AdoMet-dependent MTases^62–65^. While AdoMet binds within a buried cavity connected to the substrate binding site by a narrow channel in PCMT1 (Sup. Fig. 7), that narrow channel is blocked by a threonine to histidine substitution in PCMTD1 (Fig. 5). Selection for that histidine appears facilitated by a salt linkage to the alpha-carboxyl group of the isoaspartyl residue and the blockage of the carboxylate access to the methyl group of AdoMet. The substrate channel to this site in PCMT1 is restricted by α-helix 9, while in PCMTD1, a protruding loop increases accessibility to the binding pocket (Fig. 5B).

**Figure 5.**
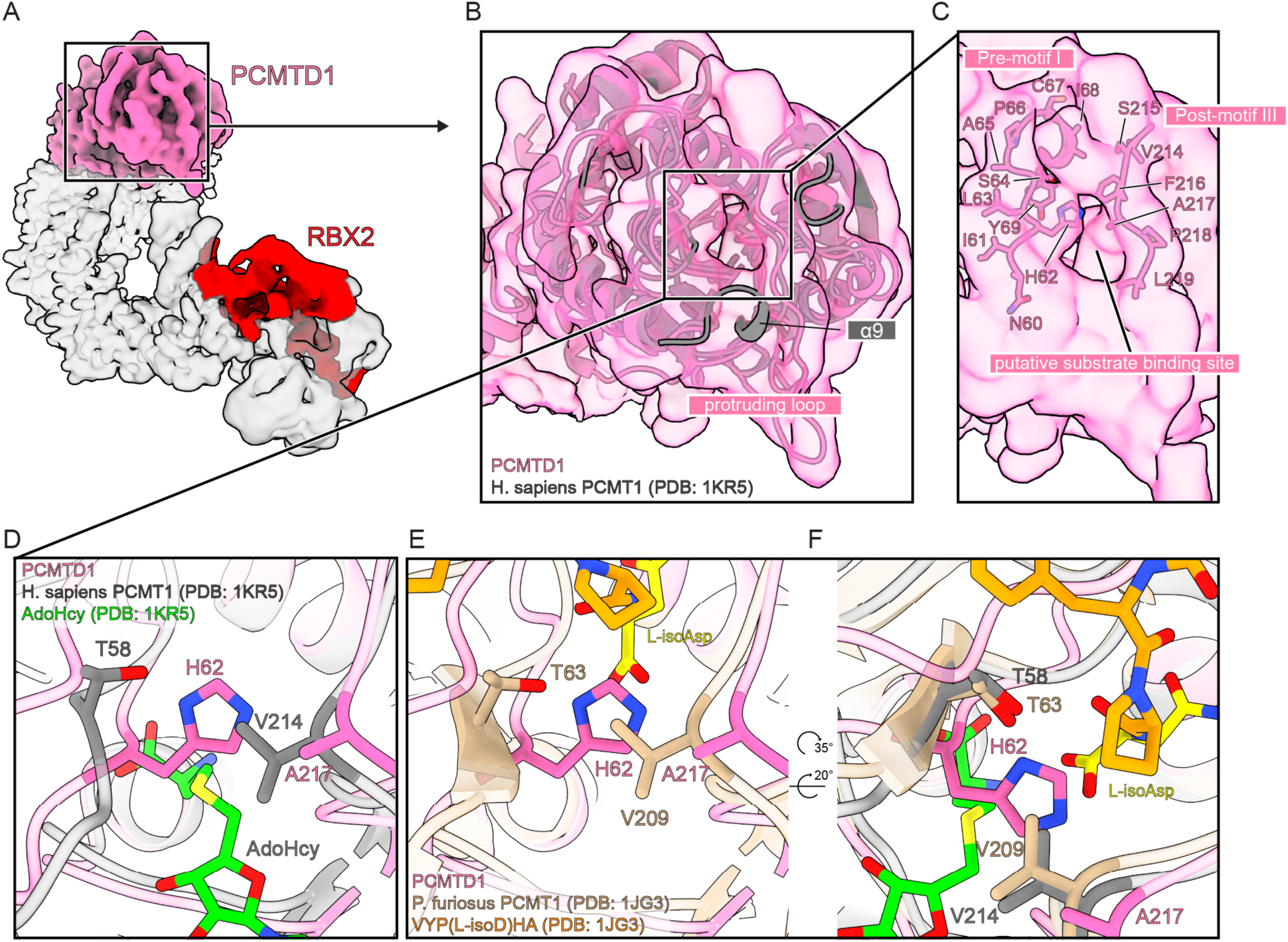
The PCMTD1 putative substrate binding site. A) Electron density for PCMTD1 and RBX2 highlighted in the CRL5-PCMTD1 complex. B) Overlay of the crystal structure of human PCMT1 against the atomic model of PCMTD1 from this study. C) Focused view on the putative substrate binding site of PCMTD1. Residues shown belong to Pre-motif I and Post-motif III on PCMTD1. D) Focused view of the AdoMet:L-isoaspartyl channel with the crystal structure of *H. sapiens* PCMT1 solved with AdoHcy aligned structurally to PCMTD1. E) Focused view of the AdoMet:L-isoaspartyl channel with the crystal structure of *P. furiosus* PCMT1 solved with VYP(L-isoasp)HA aligned structurally to PCMTD1. F) Structural alignment of *H. sapiens* PCMT1 and *P. furiosus* PCMT1 against PCMTD1 with the co-factor and peptide from respective structures shown. See also Sup. Fig. 7 depicting cross-section of the co-factor and ligand binding sites for PCMTD1 and the two variants of PCMT1.

### Neddylation induces a conformational change of the CRL5-PCMTD1 complex

The architecture observed for CRL5-PCMTD1 and its ability to recognize L-isoaspartyl targets led us to explore the structural basis for activation of substrate protein ubiquitylation. We note that recently determined structures for active CRLs display a more compact architecture compared to CRL5-PCMTD1 (Sup. Fig. 8A)^38^, as does an AlphaFold model of the complex (Sup. Fig. 8A). That prompted us to investigate structural mechanisms that may allosterically promote the ubiquitylation of L-isoaspartyl-containing protein targets, focusing on changes in CRL5-PCMTD1 induced by neddylation, a post-translational modification that broadly activates CRLs for substrate ubiquitylation^66^.

We generated stable cell lines of epitope-tagged PCMTD1 to co-immunoprecipitate (co-IP) direct and indirect protein interaction partners and compared results with a mutant variant of PCMTD1 with its BC-box point mutated to disrupt association with ELOBC, PCMTD1 P243 F247. These residues were shown to form a hydrophobic interface with ELOC from our structural data and synonymous points mutations are also known to disrupt association between PCMTD2, a paralogue to PCMTD1, and CRL components (Fig. 2)^17^. LC-MS/MS identification shows enrichment of CUL5, RBX2, ELOB, ELOC, and NEDD8 with PCMTD1 (Sup. Fig. 8B). Mutating the PCMTD1 BC-box negated these enrichments implying NEDD8 co-IPs with PCMTD1 only when the CRL5-PCMTD1 complex is formed (Sup. Fig. 8B).

We then determined the structure of neddylated CRL5-PCMTD1 by single particle cryo-EM (Fig. 6). Here, we prepared a neddylated variant of CRL5-PCMTD1 (N8-CRL5-PCMTD1) by *in vitro* neddylation and gel filtration of the reaction solution (Sup. Fig. 8C). Due to a greater degree of inherent flexibility in this complex, its structure could only be resolved to 9.72 Å resolution (Fig. 6A, Sup. Fig. 9). Nonetheless, the structure of N8-CRL5-PCMTD1 revealed a horseshoe-like architecture with long-range structural differences to CRL5-PCMTD1, as indicated by comparison to the molecular model of N8-CRL5-PCMTD1 (Fig. 6B). The termini of the neddylated complex are closer, at 150 Å apart compared to its un-neddylated parent, where they are separated by 165 Å (Fig. 1C, 6A). Additionally, the SRC is skewed ∼30° clockwise to enable this more compact architecture which results in a rearrangement and compartmentalization of the SRC-CUL5 interface (Fig. 6B,C). We propose this overall conformation caused by neddylation will further promote ubiquitylation from CRL5-PCMTD1’s elongated state.

**Figure 6.**
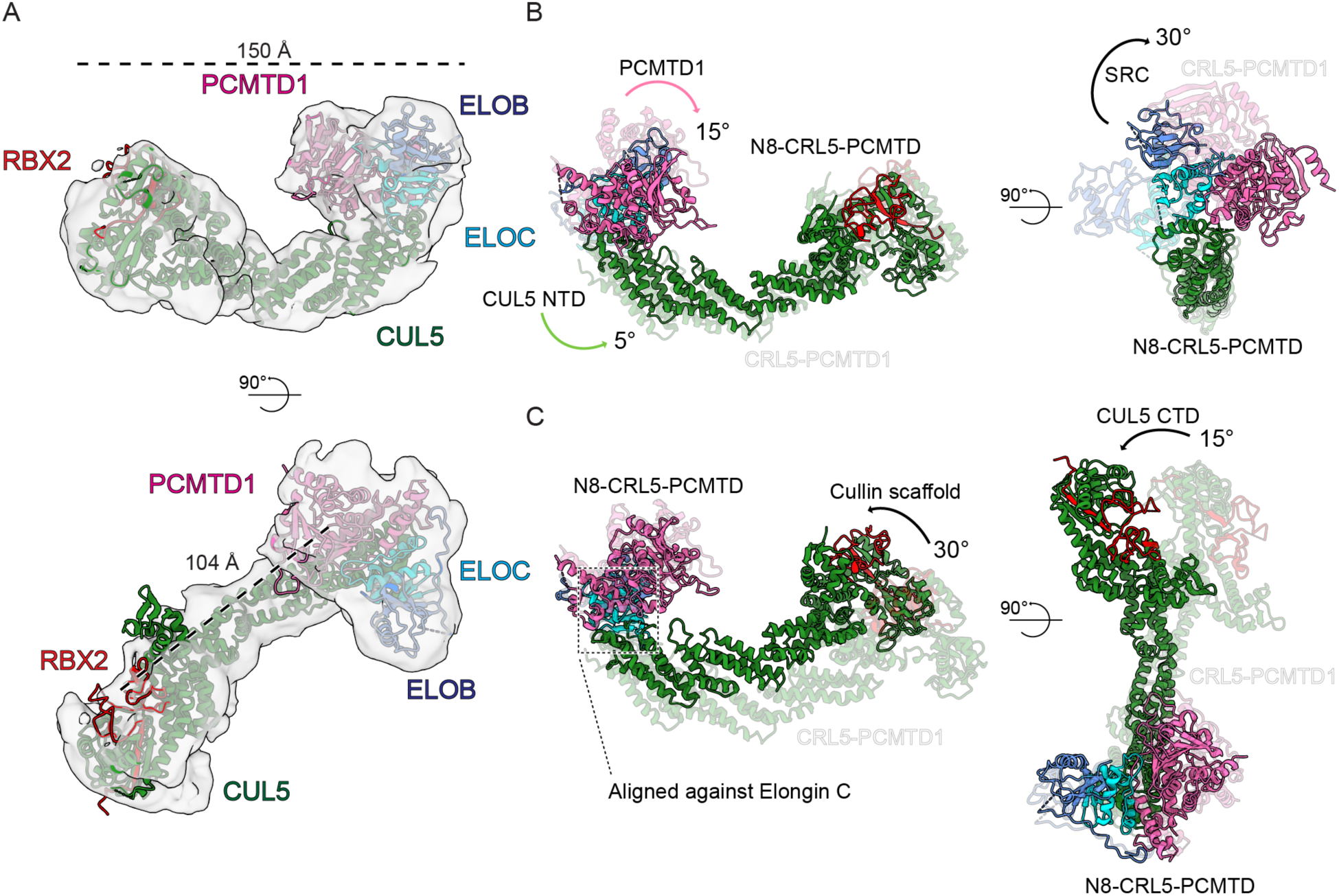
Neddylation results in a conformational change of the CRL5-PCMTD1 complex. A) Structure of CRL5-PCMTD1 after energy minimization into the cryo-EM density for neddylated CRL5-PCMTD1 with span of the complex (top) and distance between PCMTD1 and RBX2 (bottom) shown. B) The ribbon model of N8-CRL5-PCMTD1 overlayed against the structure of CRL5-PCMTD1. C) The ribbon model of N8-CRL5-PCMTD1 structurally aligned against ELOC within the structure of CRL5-PCMTD1. See Sup. Fig. 8 for sample preparation details of neddylated CRL5-PCMTD1. See also Sup. Fig. 9 for cryo-EM structural characterization details of N8-CRL5-PCMTD1.

## Discussion

To better understand regulated proteostasis of age-related spontaneous post-translational damages, we set out to define the structural basis for assembly, activation, and substrate targeting by CRL5-PCMTD1. We show that CRL5-PCMTD1 adopts a characteristic horseshoe-like architecture with conserved interfaces that stabilize the complex in a geometry reminiscent of other CRLs. PCMTD1 recognizes a specific L-isoaspartyl residue motif that is formed as a spontaneous age-related PTM, rather than an enzyme-mediated PTM or a specific amino acid sequence. Together, these findings suggest a potential new mechanism in protein turnover of age-damaged proteins via regulation through spontaneous PTMs, adding to the growing evidence for CRL recognition by substrate adaptor proteins, such as CRBN, which has been shown to recognize spontaneous C-terminal cyclic imides arising from Asn and Gln deamidation^18^.

Our structural findings for CRL5-PCMTD1 are consistent with a previously proposed mechanism for preventing the accumulation of L-isoaspartyl damaged proteins in cells, independent of the activity of PCMT1. Such a mechanism is also consistent with previous results demonstrating that knocking out PCMT1 in mice results in a limited accumulation of L-isoaspartyl levels with age, rather than a sustained increase^67^. Interestingly, higher levels of peptides containing L-isoaspartyl residues were detected in the urine of these mice, suggesting a possible proteasomal secondary mechanism for avoiding accumulation of the damaged proteins^67^. Notably, while PCMT1 knockout is embryonic lethal, its overexpression in neurons alone is sufficient to rescue that phenotype, further supporting a secondary compensatory mechanism addressing the presence of age-damaged proteins within the cell^67^.

Humans have a paralog of PCMTD1, PCMTD2^16,17^; the amino acid sequence of the two are approximately 68% identical. Thus, it is likely that PCMTD2 plays a similar physiological function as PCMTD1. On this basis, we propose that PCMT1, PCMTD1, and potentially PCMTD2, serve complementary roles for L-isoaspartyl maintenance. If so, they may target distinct subsets of proteins to mitigate the accumulation of L-isoaspartyl damage. Distinctions between the substrate binding sites of PCMTD1 and PCMT1 offer a potential basis for that proposed differential substrate targeting. While the N-terminal domain of PCMTD1 adopts a canonical 7βS-MTase fold, it also contains unique motifs absent in PCMT1, such as its protruding loop and a Met-Cys-Met cluster. Additionally, the extended SOCS-box domain of PCMTD1 features a low complexity domain that was not resolved in our cryo-EM map and has not been thoroughly characterized. These structural elements may contribute to the role of PCMTD1 in targeting specific protein substrates or enact an undiscovered alternative function. Future work is needed to fully understand their functions of these uncharacterized motifs.

Regulation of CRL5-PCMTD1 activity remains unclear. While neddylation induces long-range conformational rearrangements in the complex, the potential for local regulation of substrate engagement through AdoMet binding is intriguing as a link between metabolism and the UPS. We have shown that engagement of L-isoaspartyl-containing proteins is dependent on AdoMet binding and independently shown *in cell* and *in vitro* evidence for the neddylation-based regulation. The combination of these findings may provide evidence for potential crosstalk yielding highly specific regulation of PCMTD1’s activity. Given the dysregulation of AdoMet levels and neddylation are associated with disease, understanding how these mechanisms interplay could offer valuable insights on their roles in protein homeostasis^68,69^. Furthermore, it remains to be seen whether other mechanisms exist for L-isoaspartyl regulation, and whether other E3 Ub ligases may complement PCMTD1 in targeting age-related spontaneous protein damages. In conclusion, our work suggests that the L-isoaspartyl motif may represent a viable degron accessible by PCMTD1 and possibly PCMTD2.

## Acknowledgements

Research reported in this publication was supported as part of STROBE, an NSF Science and Technology Center through Grant DMR-1548924, and by the US National Institutes of Health (NIH) grants R35GM128867 (J.A.R.) and R35GM145286 (J.A.L.), the US National Science Foundation grant MCB-1714569 (S.G.C.), and the US Department of Energy grant DE-FC02-02ER63421 (J.A.R. and J.A.L.). Vitrification, screening, and high-resolution cryo-EM data collection was performed at the Stanford-SLAC Cryo-EM Center (S^2^C^2^), which is supported by the National Institutes of Health Common Fund Transformative High-Resolution Cryo-Electron Microscopy program (U24 GM129541). E.Z.P. was supported by the NIH National Institute of General Medical Sciences (NIGMS) predoctoral fellowship (T32GM136614). C.W.F. was supported by the NIH NIGMS predoctoral fellowship (T32GM145388). J.A.R. was also supported as a Pew Scholar and a Packard Fellow. The content is solely the responsibility of the authors and does not necessarily represent the official views of the National Institutes of Health. E.O. acknowledges support from the Undergraduate Research Scholars Program at UCLA. J.B.W. acknowledges support from the Daniel E. Atkinson and Charles A. West Undergraduate Research Fellowship at UCLA.

## Author contributions

E. Z. P., J. A. R., and S. G. C. conceptualization; E. Z. P., B. Z., J. W., J. A. L., J. A. R., and S. G. C. methodology; E. Z. P., B. Z., C. F., E. O., J. B. W., and V. P. investigation; E. Z. P., B. Z., C. F., E. O., J. B. W., V. P., J. W., J. A. L., J. A. R., and S. G. C. validation; E. Z. P., B. Z., and C. F. formal analysis; E. Z. P. visualization; J. W., J. A. L., J. A. R., and S. G. C. supervision; E. Z. P., J. A. R., and S. G. C. writing – original draft; E. Z. P., B. Z., C. F., E. O., J. B. W., V. P., J. W., J. A. L., J. A. R., and S. G. C. writing – review and editing; J. W., J. A. L., J. A. R., and S. G. C. funding acquisition; J. A. R., and S. G. C. project administration; J. W., J. A. L., J. A. R., and S. G. C. resources.

## Declaration of interests

J.A.R. is an equity stake holder of MedStruc Inc. All other authors declare no conflicts of interest.

### Supplemental material

Document S1.

Figures S1-S9

## Methods

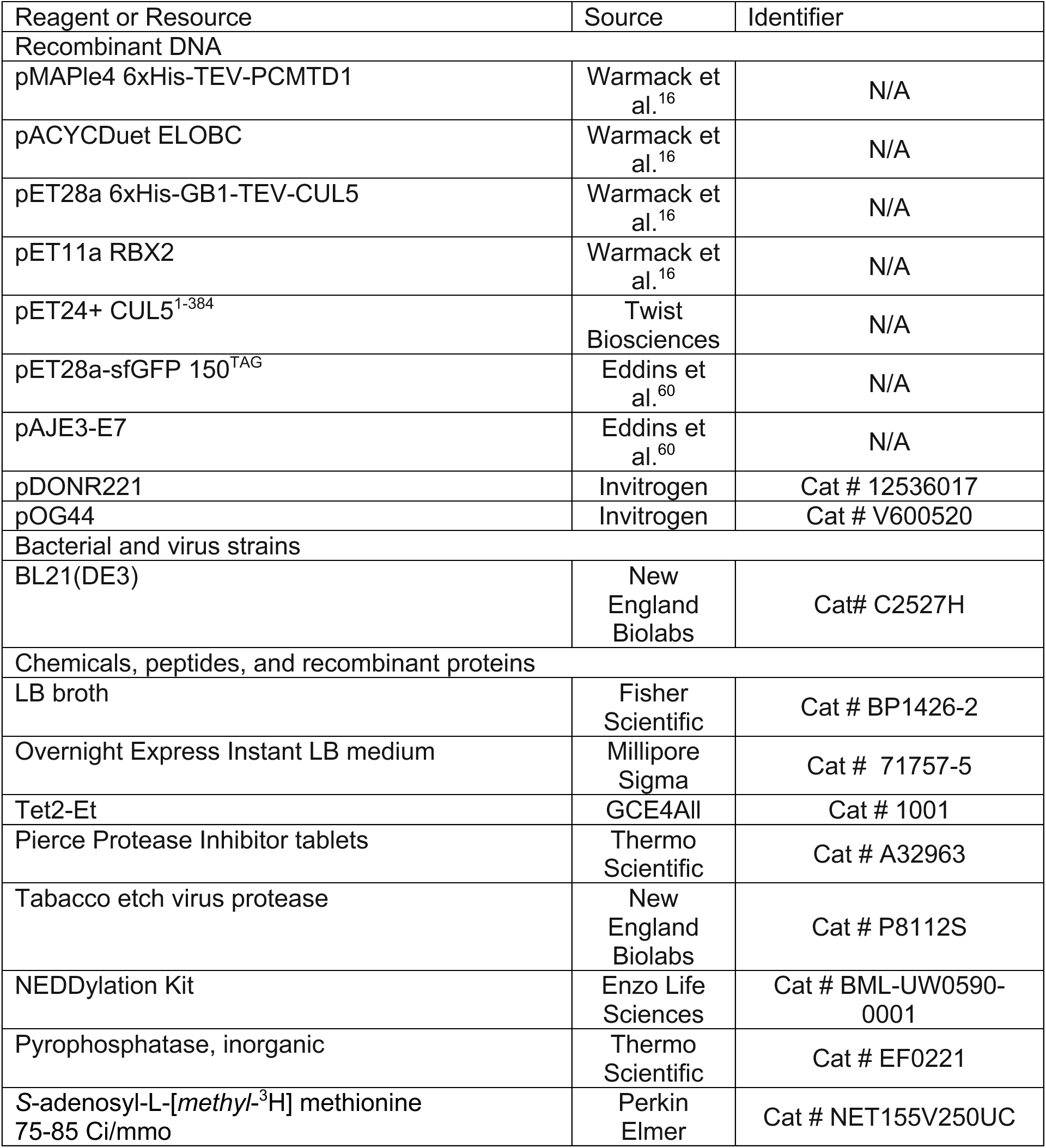
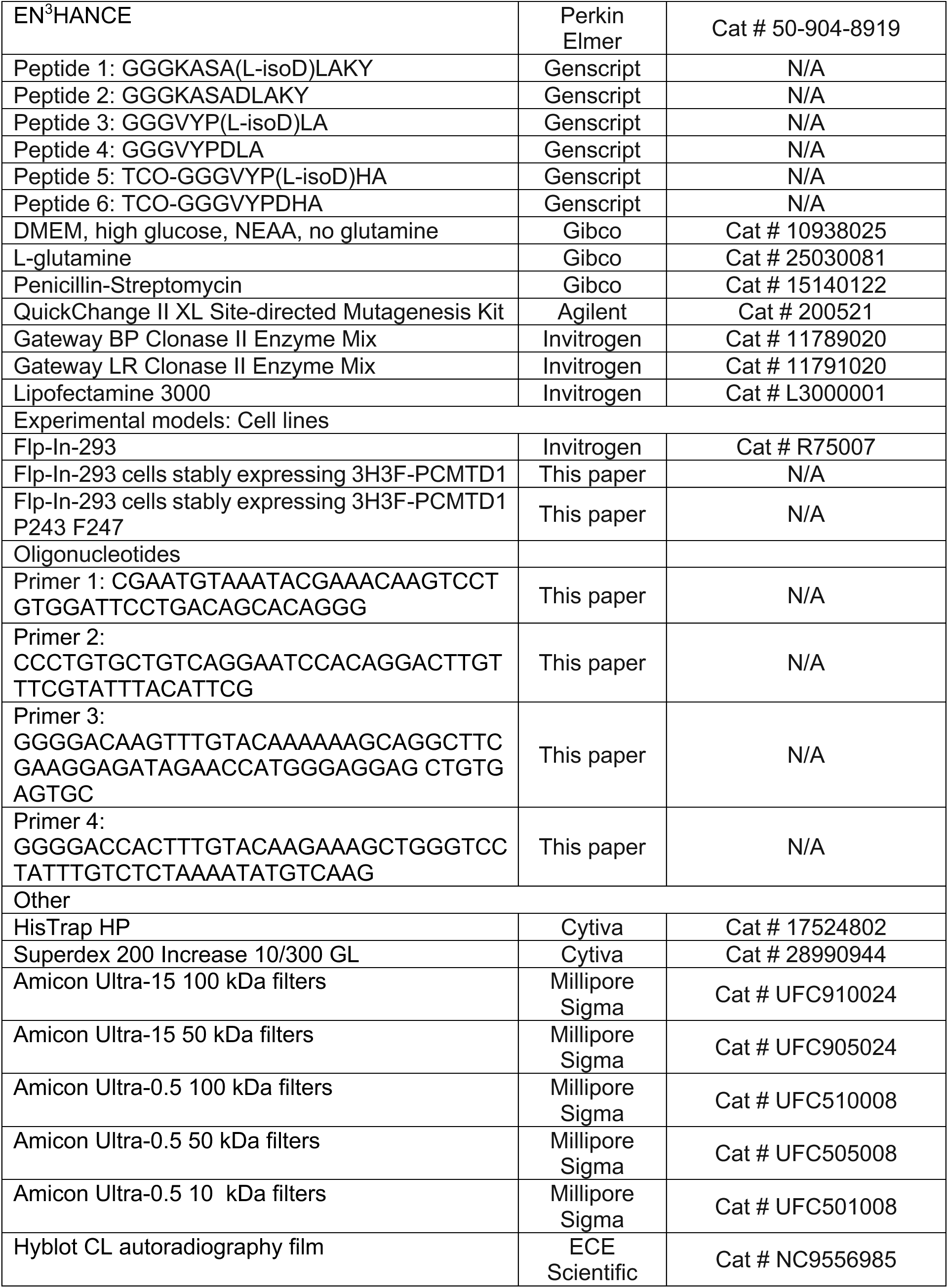
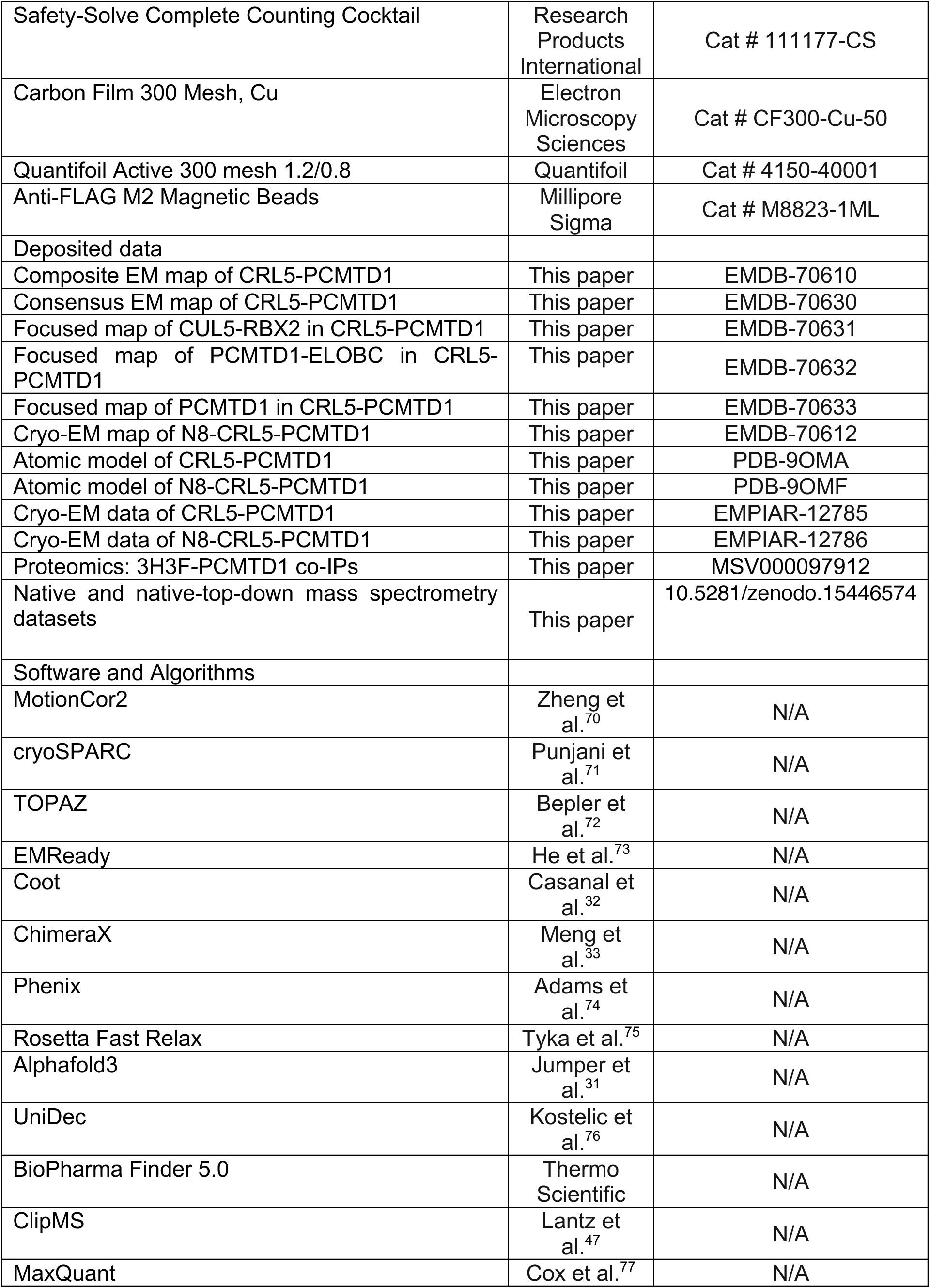

## Resource availability

### Lead contact

Further information and requests for resources and reagents generated in this study should be directed to the lead contact, Steven G. Clarke

### Materials availability

All unique cell lines and reagents generated in this study are available from the lead contact upon request as long as stocks remain available.

### Data and code availability

- All data are available in the main text or the supplemental information. Source data are provided in this paper.
- Native and native top-down mass spectrometry data are publicly accessible via Zenodo with the dataset identifier 10.5281/zenodo.15446574. Proteomic mass spectrometry data are publicly accessible via the MassIVE repository with the dataset identifier MSV000097912.
- Cryo-EM models generated in this study are publicly available as of the date of publication in the Protein DataBank via PDB entries: 9OMA for CRL5-PCMTD1 and 9OMF for N8-CRL5-PCMTD1. Corresponding density maps are available in the Electron Microscopy Data Bank, including the composite map of CRL5–PCMTD1 (EMD-70610), focused maps refined on CUL5–RBX2 (EMD-70631), PCMTD1–ELOBC (EMD-70632), and PCMTD1 (EMD-70633), the consensus map of CRL5–PCMTD1 (EMD-70630), and the EM map of N8–CRL5–PCMTD1 (EMD-70612). Raw data are publicly accessible in the Electron Microscopy Public Image Archive (EMPIAR-12785, CRL5-PCMTD1; EMPIAR-12786, N8-CRL5-PCMTD1).
- This paper does not report original code.
- Any additional information required to reanalyze the data reported in this paper is available from the lead contact upon request.

## Method Details

### Recombinant protein expression, purification, and sample preparation

Plasmids used for the recombinant expression of human 6xHis-TEV-PCMTD1 (pMAPle4), human ELOBC (pACYCDuet), human 6xHis-GB1-TEV-CUL5 (pET28a), and mouse RBX2 (pET11a) were prepared as previously described^16^. CUL5^1-384^ (pET24+) was obtained from Twist Biosciences. Plasmids used for non-canonical amino acid incorporation into sfGFP (sfGFP-Tet2) were generous gifts from Dr. Ryan Mehl and the GCE4ALL Center at Oregon State University (pET28a-sfGFP 150^TAG^ Addgene #85493, pAJE3-E7 Addgene #214359). The protocol for overexpression and purification of sfGFP-Tet2 was adapted from previous studies^60^.

Plasmids for PCMTD1 and ELOBC, CUL5 and RBX2, CUL5^1-384^, and sfGFP 150^TAG^ and the Tet2-Et tRNA synthetase/tRNA pair were co-transformed and expressed in *Escherichia coli* BL21(DE3) cells (New England BioLabs) with appropriate antibiotics. Cultures were grown in LB-media and induced at OD600 = 0.5 – 0.7 with 0.5 mM IPTG for 3 hours at 37°C for cultures containing PCMTD1 and 1.0 mM IPTG for 16 – 18 hours at 18°C for cultures containing CUL5. Cultures containing sfGFP 150^TAG^ were grown and auto-induced in Overnight Express Instant LB autoinduction media (Novagen) with 0.5 mM (S)-2-amino-3-(4-(6-ethyl-1,2,4,5-tetrazin-3-yl)phenyl) propanoic acid) (Tet2-Et; Oregon State University) for incorporation of Tet2-Et into sfGFP (sfGFP-Tet2). Following induction, all cultures were pelleted and resuspended in lysis buffer (50 mM sodium HEPES pH 7.4, 300 mM NaCl, 50 mM imidazole, 1 mM DTT, 1 tablet of Pierce protease inhibitor; Thermo Scientific), lysed by sonication, and clarified by centrifugation at 13,000 rpm for 50 min. To purify PCMTD1 in a CRL complex or as PCMTD1-CUL5^tet^, lysates for PCMTD1-ELOBC and CUL5-RBX2 or CUL5^1-384^ were combined and incubated overnight at 4°C. For all protein preparations, lysates were loaded onto 5 mL HisTrap HP columns (Cytivia) and washed (50 mM sodium HEPES pH 7.4 150 mM NaCl 50 mM imidazole 1 mM DTT) for 20 column volumes prior to elution with a linear gradient between wash buffer and elution buffer (50 mM sodium HEPES pH 7.4 150 mM NaCl 500 mM imidazole 1 mM DTT) over 5 column volumes. Fractions containing proteins were pooled, concentrated, flash frozen, and stored at -80°C.

Prior to cryo-EM or nMS analyses, all protein preparations except sfGFP-Tet2 were digested overnight with tobacco etch virus protease (New England Biolabs) at 37°C and then further purified by gel filtration (GF) using a Superdex 200 10/300 GL column (Cytivia) with GF buffer (50 mM sodium HEPES pH 7.4, 150 mM NaCl, 1 mM DTT). Fractions containing proteins were collected and concentrated with 50 or 100 kDa Millipore Amicon Ultra filters (Millapore Sigma) and used freshly.

### *In vitro* neddylation of CRL5-PCMTD1

*In vitro* neddylation of CRL5-PCMTD1 was performed using a neddylation kit according to manufacturer’s specifications (Enzo Life Sciences). Briefly, 1.5 mg of purified CRL5-PCMTD1 was incubated overnight at 37°C with a solution containing NAE1, UbcH12, NEDD8, Mg-ATP, 10 U/mL inorganic pyrophosphatase (Millapore Sigma), and 600 μM DTT. Neddylated CRL5-PCMTD1 (N8-CRL5-PCMTD1) was further enriched by gel filtration using a Superdex 200 10/300 GL column (Cytivia) with GF buffer and confirmed with SDS-PAGE and immunoblotting,

### 3H-AdoMet:protein ultraviolet light cross-linking

This protocol was adapted from previous studies^16^. In a final volume of 30 μL, proteins at a final concentration of 3.85 μM were combined with 1 μL of 0.55 mCi/mL *S*-adenosyl-L-[*methyl*-^3^H] methionine, 75-85 Ci/mmol (^3^H-AdoMet; PerkinElmer Life Sciences) in 50 mM Tris-HCl pH = 6.8 with or without excess AdoHcy or ATP at a final concentration of 0.5 mM. Reactions were then irradiated with 254 nm light for 1 hour at 4°C as described previously ^16^. After irradiation, reactions were quenched with SDS-PAGE sample buffer and electrophoresed. Following staining and de-staining, gels were then incubated with EN^3^HANCE (PerkinElmer Life Sciences), washed with water, and exposed to film (Hyblot CL; Denville Scientific) for 1 week at -80°C before development.

### Methanol vapor diffusion assay for protein ^3^H-methyl esters

This protocol was adapted from previous studies^16^. In a final volume of 100 μL with 200 mM bis-Tris-HCl pH = 6.4, 25 pmol of KASA(L-isoaspartyl)LAKY or 300 pmol of ovalbumin was combined with 100 pmol of protein and 550 pmol of ^3^H-AdoMet and incubated for 2h at 37°C. The reaction was quenched with 10 μL of 2 M sodium hydroxide and reactions were spotted on filter paper (Bio-Rad). Filter papers were then wedged in the necks of 20 mL scintillation vials pre-filled with 5 mL Safety Solve scintillation reagent (Research Products International), tightly capped, and incubated for 2 hours at room temperature. After incubation, filter papers were removed and vials were counted thrice, each at 5 min intervals, using a Beckman LS6500 scintillation counter.

### Negative-stain electron microscopy

Samples were diluted to 10 μg/mL and were deposited onto glow-discharged 300 mesh carbon film copper grids (Electron Microscopy Sciences). Grids were then stained with 1% uranyl acetate. Imaging was performed on a Tecnai T12 TEM operating at 120 keV.

### Cryo-EM sample preparation and data acquisition

Cryo-EM sample preparation and data collection was performed at the SLAC-Stanford CryoEM Center. Grid vitrification was performed using an SPT Labtech chameleon and self-wicking chameleon grids (Quantifoil Active 300 mesh 1.2/0.8; Quantifoil). Prior to grid vitrification, samples were freshly purified and diluted to 2.8 mg mL^-1^ and 2.3 mg mL^-1^ for CRL5-PCMTD1 and N8-CRL5-PCMTD1, respectively, in GF buffer. 7 uL of purified samples were loaded onto the chameleon system. A total of 40 nL of sample was dispensed onto glow discharged grids and plunge frozen into liquid ethane. After visual inspection of wicked grids, accepted grids were stored in liquid nitrogen until screening and data collection.

Cryo-EM data collection was performed using a Titan Krios G3 (ThermoFisher Scientific) in counting mode operating at 300 keV and equipped with a BioQuantum K3 energy filter (Gatan) for CRL5-PCMTD1 and a Titan Krios G3i (ThermoFisher Scientific) in counting mode operating at 300 keV and equipped with a Selectris X imaging filter (Thermo Scientific) for N8-CRL5-PCMTD1. With a total dose of 50 e^-^/Å^2^, data were collected at 0.86 Å per pixel on the Titan Krios G3 and 0.743 Å per pixel on the Titan Krios G3i . The nominal defocus range used for data collection was -1.5 to -2.5 μm. Detailed collection parameters are summarized in Table 1.

**Table 1.** Cryo-EM data collection, refinement and validation statistics. ^a^ = this study

|  | CRL5-PCMTD1 | N8-CRL5-PCMTD1 |
| --- | --- | --- |
| Data collection |  |  |
| Microscope | Krios G3 | Krios G3i |
| Voltage (kV) | 300 | 300 |
| Total electron exposure (e<sup>-</sup>/Å<sup>2</sup>) | 50 | 50 |
| Defocus range (μm) | ~-1.5 – -2.5 | ~-1.5 - -2.5 |
| Pixel size (Å) | 0.86 | 0.743 |
| Micrographs collected | 5,447 | 16,925 |
| Micrographs used | 2,773 | 12,280 |
| Reconstruction |  |  |
| 3D processing package | cryoSparc | cryoSparc |
| Initial particle images (no.) | 1,302,046 | 6,609,912 |
| Final particle images (no.) | 352,937 | 7,521 |
| Symmetry | C1 | C1 |
| FSC 0.143 (unmasked) | 4.14 | 9.72 |
| Local resolution range (Å) | - | - |
| Map sharpening package | EMReady | - |
| Refinement |  |  |
| Initial model used (PDB code) | 6V9I, 8FVI, Alphafold | 9OMA<sup>a</sup> |
| Model refinement package | Rosetta Fast Relax, Phenix | Rosetta Fast Relax, Phenix |
| Nonhydrogen atoms | 10,328 | 10,303 |
| B factor (Å<sup>2</sup>) | - | - |
| Protein residues | 1,270 | 1,267 |
| RMS deviations | 0 | 0 |
| Validation |  |  |
| ClashScore | 8 | 5 |
| Poor rotamers | 2% | 1% |
| Ramachandran plot |  |  |
| Favored (%) | 94.14% | 94.93% |
| Allowed (%) | 5.46% | 4.75% |
| Disallowed (%) | 0.40% | 0.32% |

<sup>a</sup> = this study

### Image processing and model building

A total of 5,447 and 15,192 micrographs were collected for CRL5-PCMTD1 and N8-CRL5-PCMTD1, respectively, and were aligned and dose-weighed with MotionCor2^70^ for the former dataset and cryoSPARC Patch Motion Correction^71^ for the latter dataset. Micrographs with a reported CTF resolution ≥ 4.5 Å were discarded for CRL5-PCMTD1 leaving 2,773 micrographs. Similarly, micrographs with a reported CTF resolution ≥ 5 Å for N8-CRL5-PCMTD1 were discarded leaving 12,280 micrographs. Subsequent data processing was performed using cryoSPARC versions 4.1.0-4.3.0 and the processing pipeline is summarized in Sup. Fig. 2 and detailed further below^71^.

For CRL5-PCMTD1, blob picking was used to extract 1,302,046 particles (4x binned) for initial 2D classification. 2D classes containing strong secondary features were subjugated to ab initio reconstruction (n = 3) which generated an initial model used for further 3D refinement. These classes were also used as templates for template picking and an additional 861,626 particles were picked and combined with the initial particle set. After removing duplicate particles, the resulting 1,487,781 particles were then subjugated to six rounds of heterogenous refinement using the initial 3D model as reference and five decoy classes that were generated from 2D classes of obvious junk particles. After heterogeneous refinement, the resulting 214,183 particles were re-extracted to the full pixel size. These particles were then supplemented with Topaz-picked particles (144,361) trained with 2D templates of under-represented classes in the data set^72^. Subsequently, homogeneous refinement, particle-recentering to the SRC, and non-uniform refinement was used to generate an initial consensus map of the complex. Density and features for the region containing PCMTD1 were low-resolution in the consensus map. Thus, further local classification was implemented using masks encompassing either PCMTD1, the SRC, or the

Cullin-scaffold to obtain higher-resolution final maps. Local and global CTF refinement was also tested but yielded no improvement. The final maps were then sharpened with EMReady (beta version 2.0)^73^ for model building and resolutions of unsharpened maps were reported based on the gold-standard Fourier shell correlation 0.143 criterion.

Model building using sharpened maps was performed using Coot and UCSF ChimeraX^32,33^. Visualization, evaluation of 3D density maps, and generating figures were performed with UCSF ChimeraX^33^. To build the model of CRL5-PCMTD1, a starting model of the complex was generated by 1) docking the published cryo-EM structure of CUL5-RBX2 (PDB 6V9I) into the consensus map after protein side-chain packing with FASPR^78^, 2) fitting the crystal structure of ELOBC by structurally aligning CUL5 from the published model of APOBEC3H-Vif-CBF-beta-EloB-EloC-CUL5 (PDB 8FVI) to the docked model of CUL5-RBX2, and 3) positioning the predicted model of PCMTD1 in the map by sequence alignment against the SOCS-box of Vif all within UCSF ChimeraX^30,55^. Following energy minimization in the cryo-EM map with FastRelax, the starting model of the complex was then iteratively adjusted into cryo-EM maps with Coot and refined with PHENIX’s phenix_real_space_refine including H-bond restraints generated from highly-confident secondary elements present in the predicted model of PCMTD1^32,74,75^.

For N8-CRL5-PCMTD1, micrographs were first denoised prior to blob picking in which 1,736,071 particles (4x binned) were extracted for initial 2D classification. 2D classes containing strong secondary features were subjugated to ab initio reconstruction (n = 4) which generated an initial model used for further 3D refinement. These classes were also used as templates for initial template picking and an additional 1,995,353 particles were picked and combined with the initial particle set. After removing duplicate particles, the resulting 2,886,961 particles were then subjugated to four rounds of heterogenous refinement using the initial 3D model as reference and five decoy classes that were generated from 2D classes of obvious junk particles. After heterogeneous refinement, the resulting 53,538 particles were re-extracted to the full pixel size followed by subsequent rounds of 3D and 2D classification to generate new templates for another round of template picking resulting in an additional 5,233,956 picked particles. After removing duplicates, the final 6,609,912 particles were subjected to 11 additional rounds of heterogeneous refinement resulting in a particle stack of 55,695 particles. After re-extraction to full box size, the particles were then surjected to 3 further rounds of 3D classification to minimize particle heterogeneity, and after non-uniform refinement, lead to the consensus map made up of 7,521 particles.

To generate the model of N8-CRL5-PCMTD1, the determined structure of CRL5-PCMTD1 from this study was energy minimized into the cryo-EM map of N8-CRL5-PCMTD1 with Rosetta FastRelax. The energy-minimized model was then iteratively assessed and adjusted into cryo-EM maps with Coot and refined with PHENIX’s phenix_real_space_refine primariliy with rigid_body fitting, secondary structure restraints generated from secondary features already present in the model of CRL5-PCMTD1, and Ramachandran restraints^32,74,75^.

### Generating sfGFP peptide conjugates

To conjugate peptides to sfGFP-Tet2, peptides with an N-terminal trans-cyclooctene (TCO) were obtained from Genscript. For each reaction, 94 μL of 16 μM sfGFP-Tet2 was combined with 15 μL of 100 μM TCO-peptide and brought to a final volume of 150 uL with PBS. The reaction was then incubated at room temperature overnight. Following the reaction, unreacted excess peptides were removed by three rounds of buffer exchanges into PBS using 10 kDa Millipore Amicon Ultra filters. Reaction completion was then assessed by nMS (Sup. Fig 6). If not used immediately, proteins were flash frozen and stored at -80°C.

### Native mass spectrometry (nMS)

#### Sample preparation

Theoretical molecular weights for protein species and peptides used in nMS experiments are listed in Table 2. Proteins were buffer exchanged with 50 kDa or 100 kDa Millipore Amicon Ultra filters into 200 mM ammonium acetate and diluted to a final concentration of 20 μM. Peptides were obtained from GenScript and dissolved in MS-grade water to a final concentration of 50 μM. AdoMet was dissolved in MS-grade water to a final concentration of 50 μM. For protein-binding experiments, proteins, peptides or AdoMet were combined at a 1:1 molar ratio just prior to analysis.

**Table 2.** Theoretical MW’s for nMS analytes. ^a,b^ = N-terminal methionine cleavage observed in experiments

| Recombinant proteins and peptide analytes | Theoretical MW (Da) |
| --- | --- |
| PCMTD1 | 40,761 |
| ELOB | 13,133 |
| ELOC | 10,964 |
| ELOBC | 24,096 |
| CUL5 | 91,288 |
| RBX2 | 12,707 |
| PCMTD1-ELOB | 53,894 |
| PCMTD1-ELOC | 51,725 |
| PCMTD1-ELOBC | 64,857 |
| CUL5-RBX2 | 103,995 |
| CUL5(1-384)<sup>a</sup> | 44,392 |
| PCMTD1- CUL5<sup>tet,b</sup> | 109,247 |
| CRL5-PCMTD1 | 168,853 |
| PCMT1 | 29,536 |
| KASA(L-isoD)LAKY | 966 |
| GGGKASADLAKY | 1,137 |
| GGGKASA(L-isoD)LAKY | 1,137 |
| GGGVYPDLA | 848 |
| GGGVYP(L-isoD)LA | 848 |
| sfGFP-Tet2 | 27,968 |
| TCO-GGGVYPDHA | 1,025 |
| TCO-GGGVYP(L-isoD)HA | 1,025 |
| sfGFP-Tet2-TCO-GGGVYPDHA | 28,969 |
| sfGFP-Tet2-TCO-GGGVYP(L-isoD)HA | 28,969 |

**Table 3.**
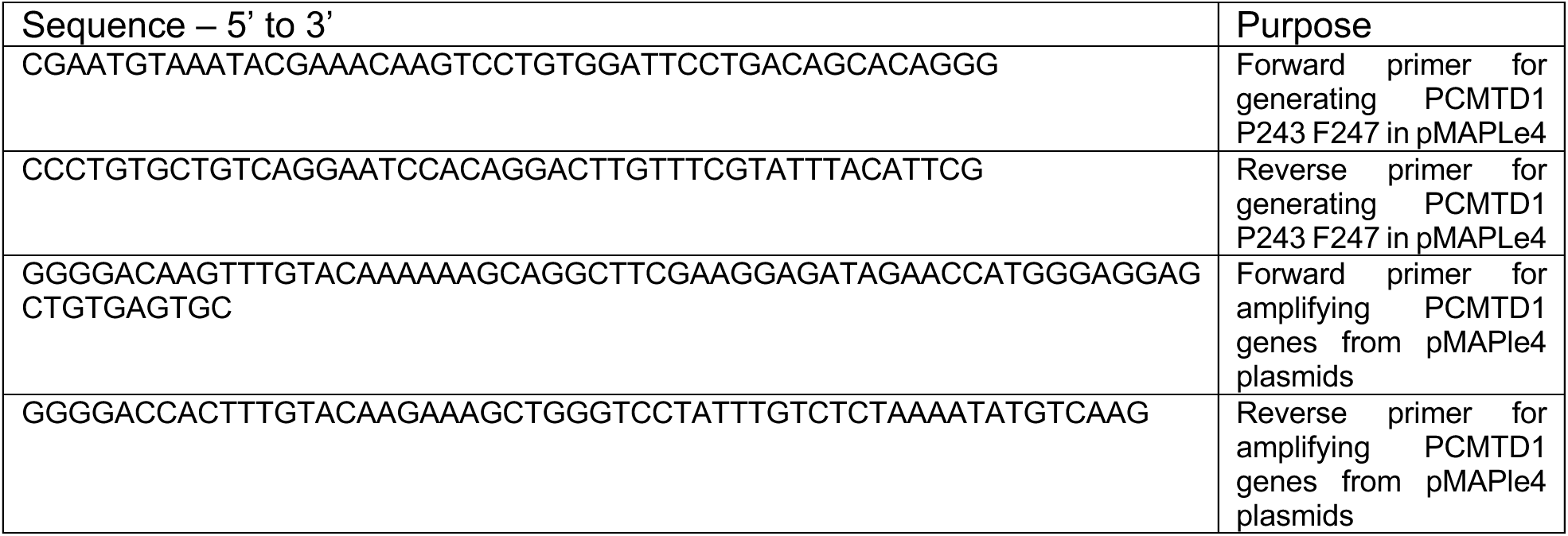
Primers used for stable cell line generation.

### Native MS and Native Top-Down MS

Native MS and HCD experiments were performed with a Thermo Fisher Scientific Q Exactive UHMR Orbitrap mass spectrometer with an ExD cell developed by e-MSion. Protein solutions were loaded into Au or Pt-coated borosilicate capillaries on a nanospray ionization source and sprayed at 0.8-1.6 kV, 175 °C with the S-lens RF set at 100 V. Native mass spectra were collected at a resolution of 12,500 at m/z 400 and all ion parameters were optimized to focus on the native protein-ligand analytes using voltage rollercoaster filtering methods^79^. In-source CID was set over a range of 30-150 eV for monomer ejection experiments. A single charge state was isolated by the quadrupole for protein complex and subcomplexes. The HCD collision energy was optimized to minimize the generation of b/y fragment ions. All top-down mass spectra were acquired at 100,000 resolution (m/z 400) with a noise threshold set to 3. Each spectrum was a result of 500 averaged scans.

### Native mass spectra data analysis

Native spectra were deconvoluted by UniDec^76^. All MS/MS spectra were deconvoluted with BioPharma Finder 5.0 (ThermoFisher Scientific, Waltham, USA). Deconvolved peaks were assigned by ClipsMS 2.0.035 with an error tolerance of 5 ppm^80^. Sequence assignment accommodated the major ECD (c, c+H, c-H, z, z+H, z-H) and HCD (b, y) ion types without annotating neutral losses ions, except when explicitly mentioned. Terminal fragments were manually validated by confirming the isotopic distributions.

### Stable cell line generation and co-immunoprecipitation experiments

#### Cell culture

Flp-In-293 cells (Thermo Fisher Scientific) and all cell lines generated from Flp-In-293 cells were cultured in Dulbecco’s Modified Eagle Medium (DMEM; Thermo fisher Scientific) supplemented with 10% fetal bovine serum and 50 U/mL penicillin-streptomycin (Thermo Fisher Scientific).

### Vector cloning

The PCMTD1 construct in pMAPle4 vector was used as a template for site directed mutagenesis (using primers 1 and 2) with the QuickChange II XL Site-Directed Mutagenesis kit (Agilent) to generate PCMTD1 P243 F247. Within the pMAPle4 plasmids, the coding region of PCMTD1 and PCMTD1 P243 F247 were then amplified with flanking att*B* sequences (using primers 3 and 4) and introduced into the pDONR221 vector (Invitrogen) with BP Clonase II (Invitrogen). After incorporation into pDONR221 vectors, the PCMTD1 genes were then swapped into pDEST-3H3F-TO with LR Clonase II (Invitrogen). The pDEST-3H3F-TO vector allows hygromycin B resistance and provides tetracycline / doxycycline inducible gene expression of incorporated genes with an N-terminal 3xHA-3xFLAG epitope tag.

### Transfections and stable cell line generation

For stable cell line incorporation, Flp-In-293 cells were then co-transfected with pOG44 (Invitrogen) and pDEST-3H3F-PCMTD1 or pDEST-3H3F-PCMTD1 P243 F247 using Lipofectamine 3000 (Invitrogen). Cells were then treated with 100 μg/mL hygromycin B for 3 weeks for positive selection. Following selection, cells with the expression vectors stably incorporated were expanded, flash frozen, and stored in liquid nitrogen until further use.

### PCMTD1 immunoprecipitation and co-immunoprecipitation of interacting proteins

PCMTD1 inducible cells and PCMTD1 P243 F247 inducible cells were expanded to twelve 15 cm^2^ plates and induced with 1 μg/mL doxycycline. After 48 hours of induction, cells were harvested and lysed with 10 mL immunoprecipitation (IP) buffer (100 mM Tris HCl pH 8.0, 150 mM NaCl, 2.5 mM EDTA, 5 mM MgCl2, 5% glycerol, 0.1% NP-40, 1 mM DTT, Pierce Protease inhibitor, 1x Phosphatase inhibitor). Lysates were then clarified by centrifugation. After transferring clarified lysates to fresh 15 mL tubes, 80 μL of anti-FLAG M2 magnetic beads (Sigma-Aldrich) was added and the lysate solutions were incubated for 2 hours at 4°C with gentle rocking. Immunoprecipitates were then washed three times with IP buffer before elution with 230 μL 8 M urea in 100 mM Tris-Cl pH 7.5. Eluates were then analyzed by SDS-PAGE, immunoblotting, and LC-MS.

### LC-MS identification of immunoprecipitated proteins

Eluates in 8 M urea and 100 mM Tris-Cl (pH 8.5), obtained after immunoprecipitation, were reduced with 5 mM tris(2-carboxyethyl)phosphine and alkylated with 10 mM iodoacetamide. The reduced and alkylated proteins were then cleaned using the single-pot, solid-phase-enhanced sample preparation (SP3) protocol for protein purification^81^. Following cleanup, proteins were digested overnight at 37 °C with the proteases Lys-C and trypsin. The resulting peptides were further purified using an offline SP3-based cleanup protocol, dried, and analyzed by LC-MS/MS. Details of the LC-MS/MS method are described in^82^. Mass spectrometry data were processed using the MaxQuant bioinformatics pipeline ^77^, with peptide identification carried out by the Andromeda search engine against the *Homo sapiens* reference proteome (UniProt ID: UP000005640). Key settings included: a maximum of two missed cleavages, a false discovery rate (FDR) of 1% for both peptides and proteins, label-free quantification (LFQ) enabled with a minimum LFQ ratio count of 1, and parent and precursor ion tolerances of 20 and 4.5 ppm, respectively. Processed MaxQuant output files were further analyzed for differential protein enrichment using the *artMS* (Analytical R tools for Mass Spectrometry) pipeline^83^.

