## Supplemental Figures for "Structural basis for L-isoaspartyl-containing protein recognition by the PCMTD1 cullin-RING E3 ubiquitin ligase"

**Title**:

**Supplementary Figures**

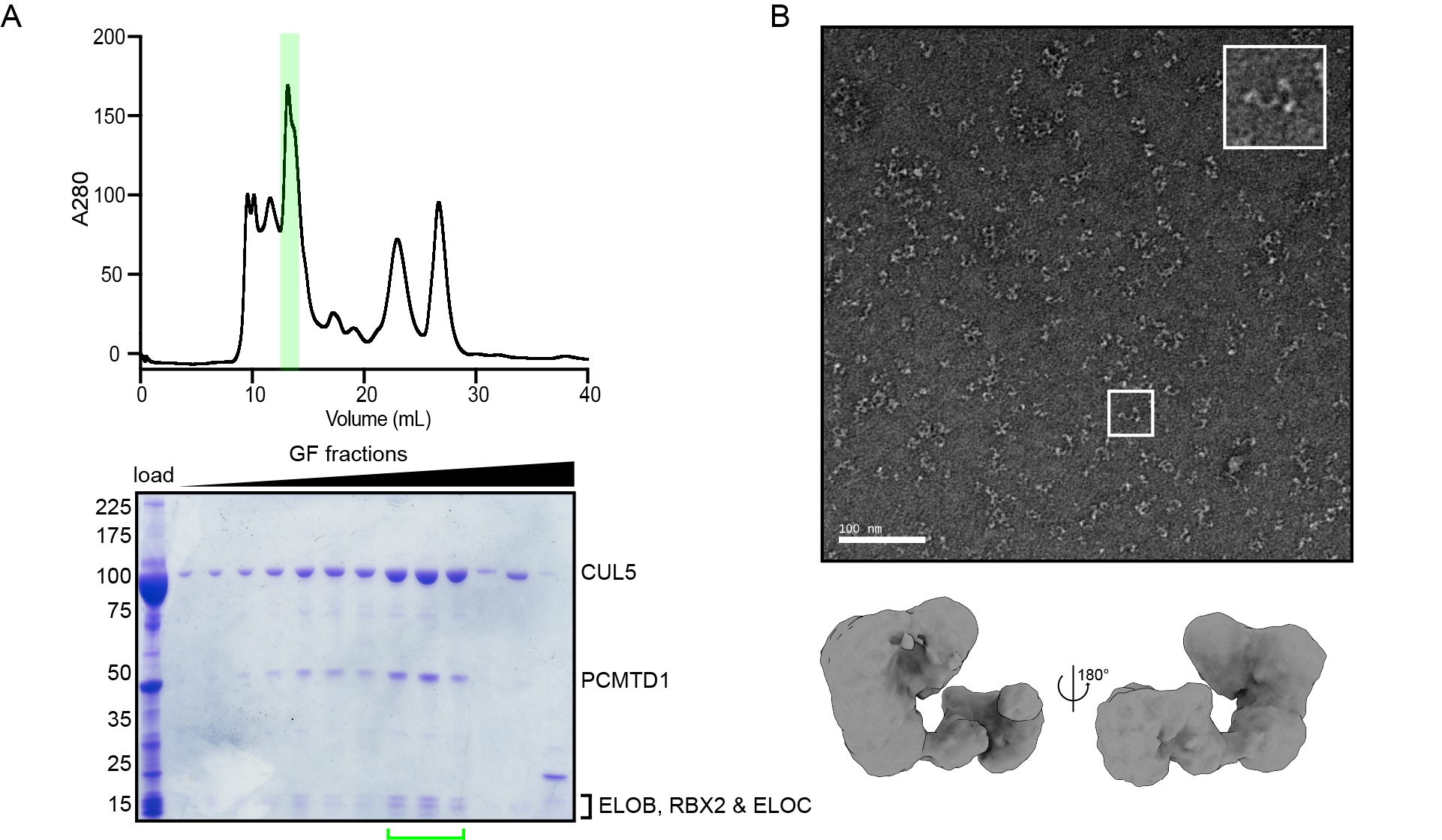

**Supplementary Figure 1. CRL5-PCMTD1 sample preparation, related to Figure 1.** A) Gel filtration chromatogram and corresponding protein gel used for enriching CRL5-PCMTD1. Green represents fractions pooled for structural and biochemical characterization. B) Negative stain electron micrographs (top) and negative stain 3D reconstruction (bottom) of CRL5-PCMTD1.

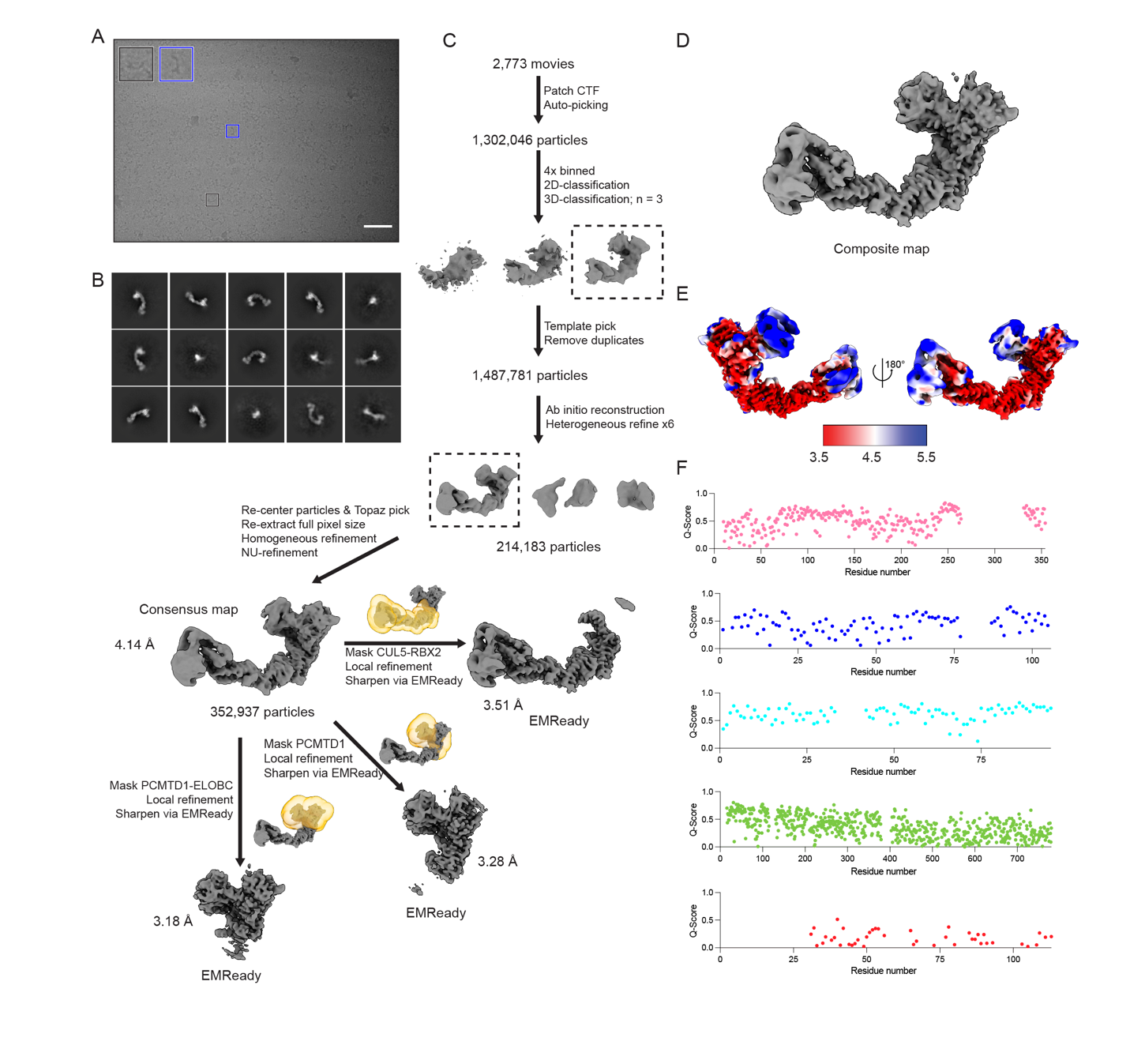

**Supplementary Figure 2. Cryo-EM 3D reconstruction pipeline for CLR5-PCMTD1, related to Figure 1.** A) Cryo-EM micrographs of CRL5-PCMTD1 after sample vitrification using the Chameleon system. Scale bar = 50 nm. B) 2D class averages of CRL5-PCMTD1 from cryo-EM datasets. C) Data processing pipeline for CRL5-PCMTD1 with masks used for local refinement. D) Composite map of CRL5-PCMTD1 following local refinement. E) Local resolution of the consensus map of CRL5-PCMTD1. Values are in angstroms. F) Q-score analysis of CRL5-PCMTD1 model-to-map fit. Pink = PCMTD1; blue = ELOB; cyan = ELOC; green = CUL5; red = RBX2.

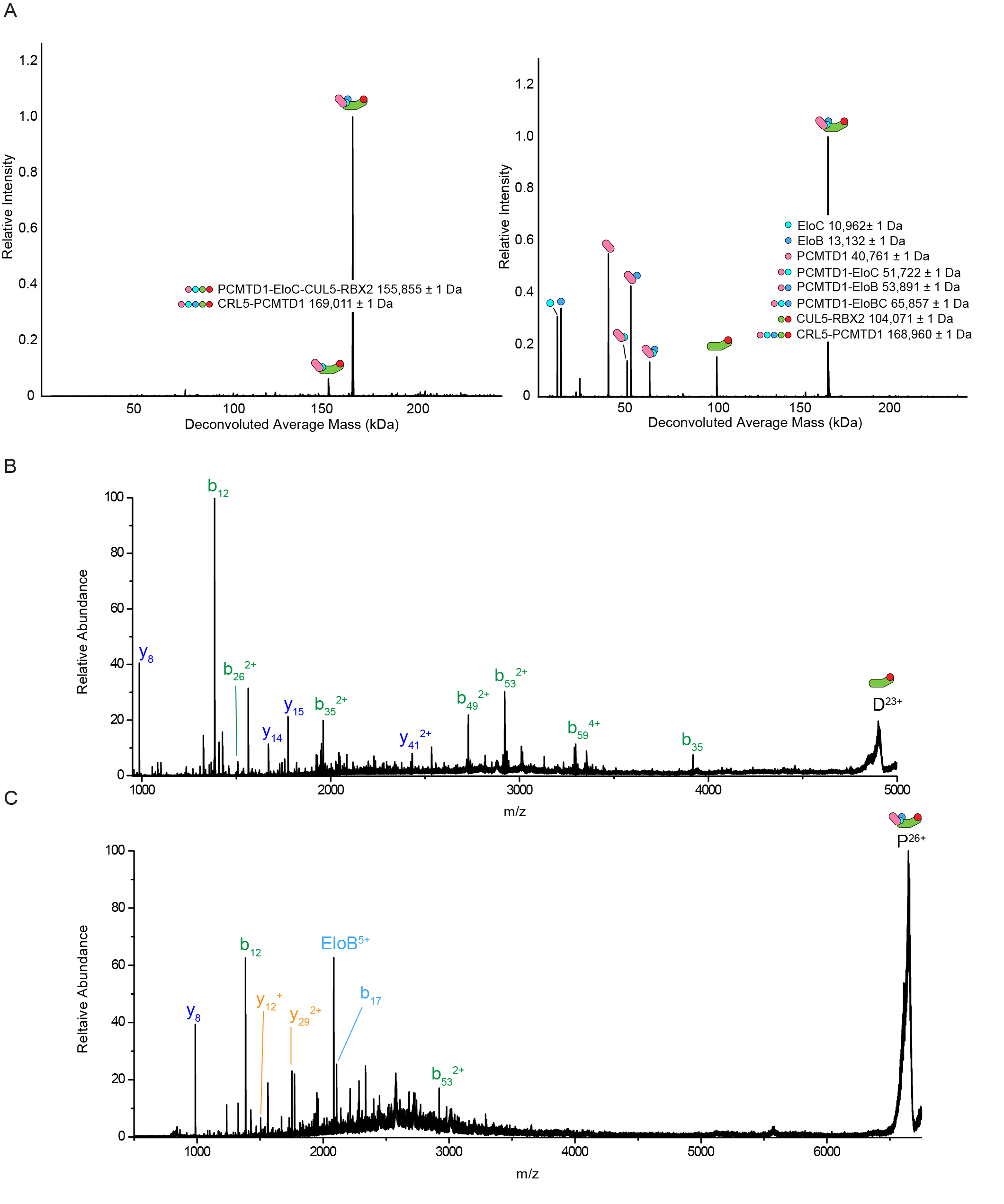

**Supplementary Figure 3.** **Native and native top-down characterization of CRL5-PCMTD1, related to Figure 1.** A) Deconvolved native mass spectra of fresh preparations of CRL5-PCMTD1 (left) and after prolonged storage in -80°C (right). B) HCD fragmentation of the CUL5-RBX2 dimer. N-terminal *b*-ions released from CUL5 are in green, and C-terminal *y*-ions from RBX2 are in blue. C) HCD fragmentation of CRL5-PCMTD1 pentamer. Ejected EloB monomer and N-terminal fragment from ELOB are in cyan; C-terminal fragments from PCMTD1 are in yellow; N-terminal fragments from CUL5 are in green; and C-terminal fragments from RBX2 are in blue.

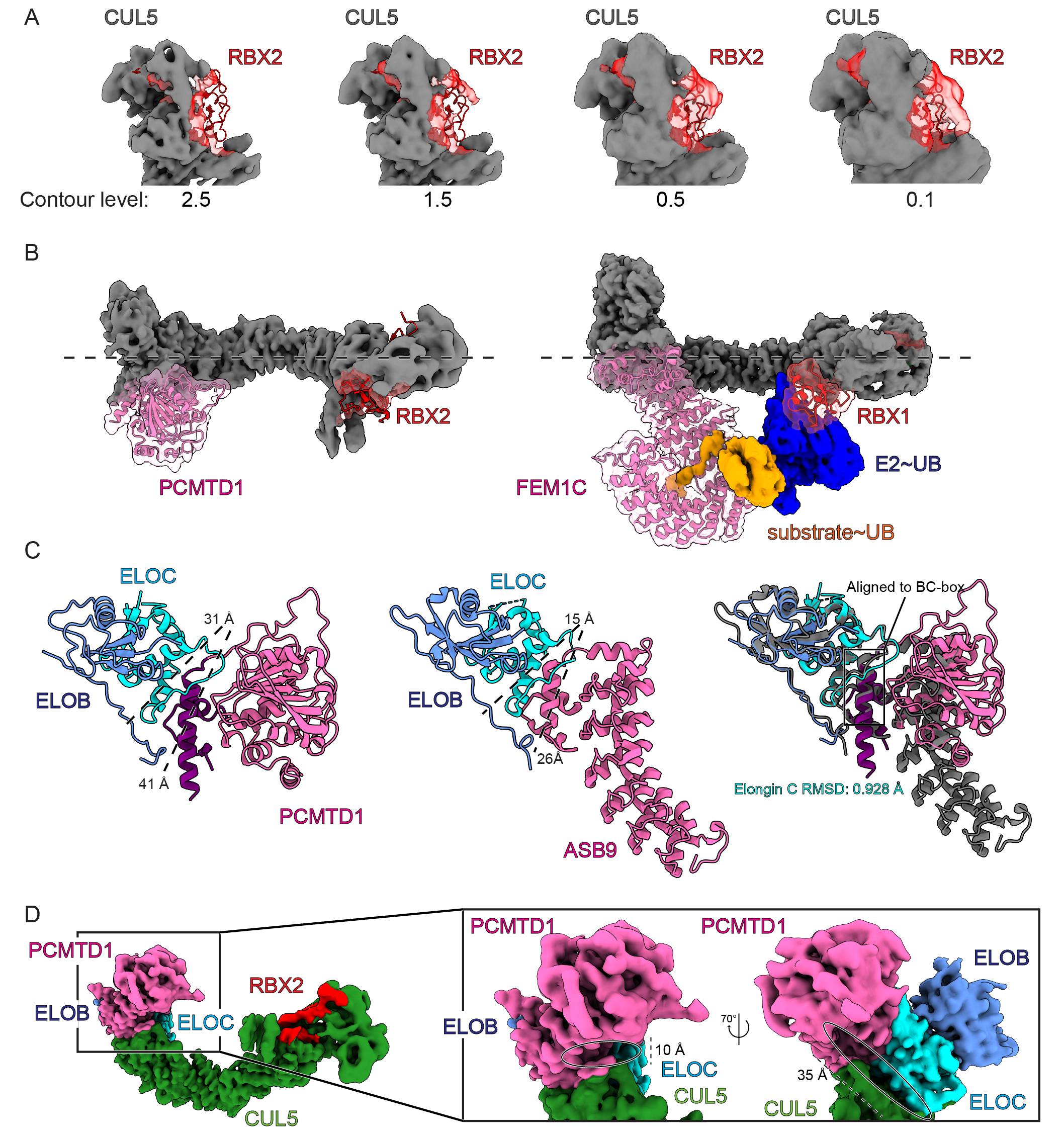

**Supplementary Figure 4. Structural features of CRL5-PCMTD1 are comparable to other CRLs, related to Figure 2.** A) Focused view of RBX2 within the cryo-EM map of CRL5-PCMTD1 at different contour levels. B) Top-down view of CRL5-PCMTD1 (left) and CRL2-FEM1C trapped in an active ubiquitylation conformation (right, PDB = 8PQL). C) Focused view of the ELOBC interface against PCMTD1 (left) and ASB9 (middle, PDB = 6V9H) followed by structural alignment against ELOC (right). D) Focused view of the groove which spans the SRC-cullin interface adjacent to the amino-terminal of the CUL5.

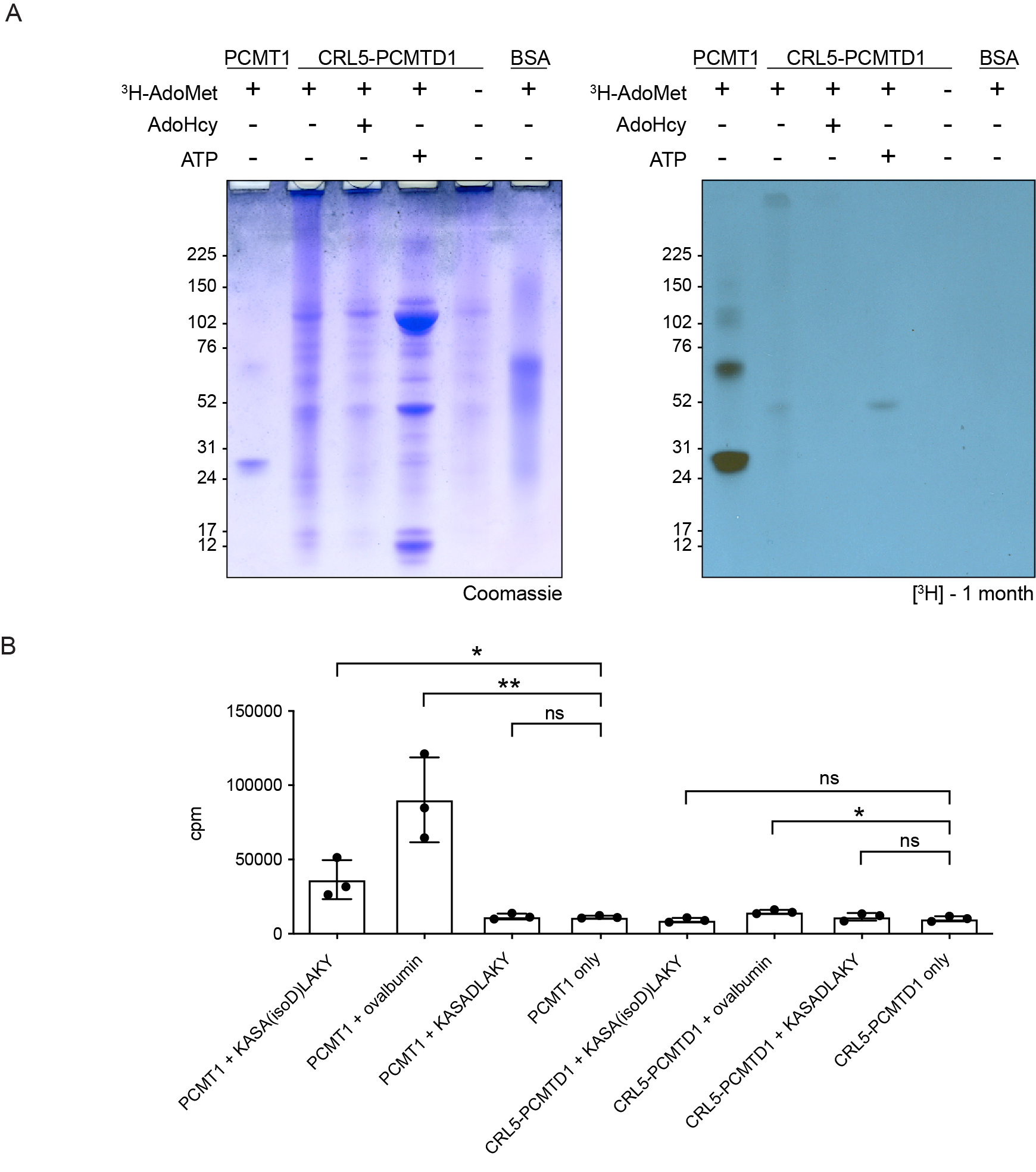

**Supplemental Figure 5. Biochemical characterization PCMTD1 when complexed as the higher-order CRL5-PCMTD1 complex, related to Figure 3.** A) 3H-AdoMet UV-crosslinking experiments with Coomassie (left) and exposed film (right) shown. Film was developed after 1 month of exposure against dried gels. B) Methanol vapor diffusion results for CRL5-PCMTD1 against substrates containing L-isoaspartyl damages. cpm = counts per minute. Error bars = standard deviation. Unpaired T-test: * < 0.05; ** < 0.005; ns = not significant.

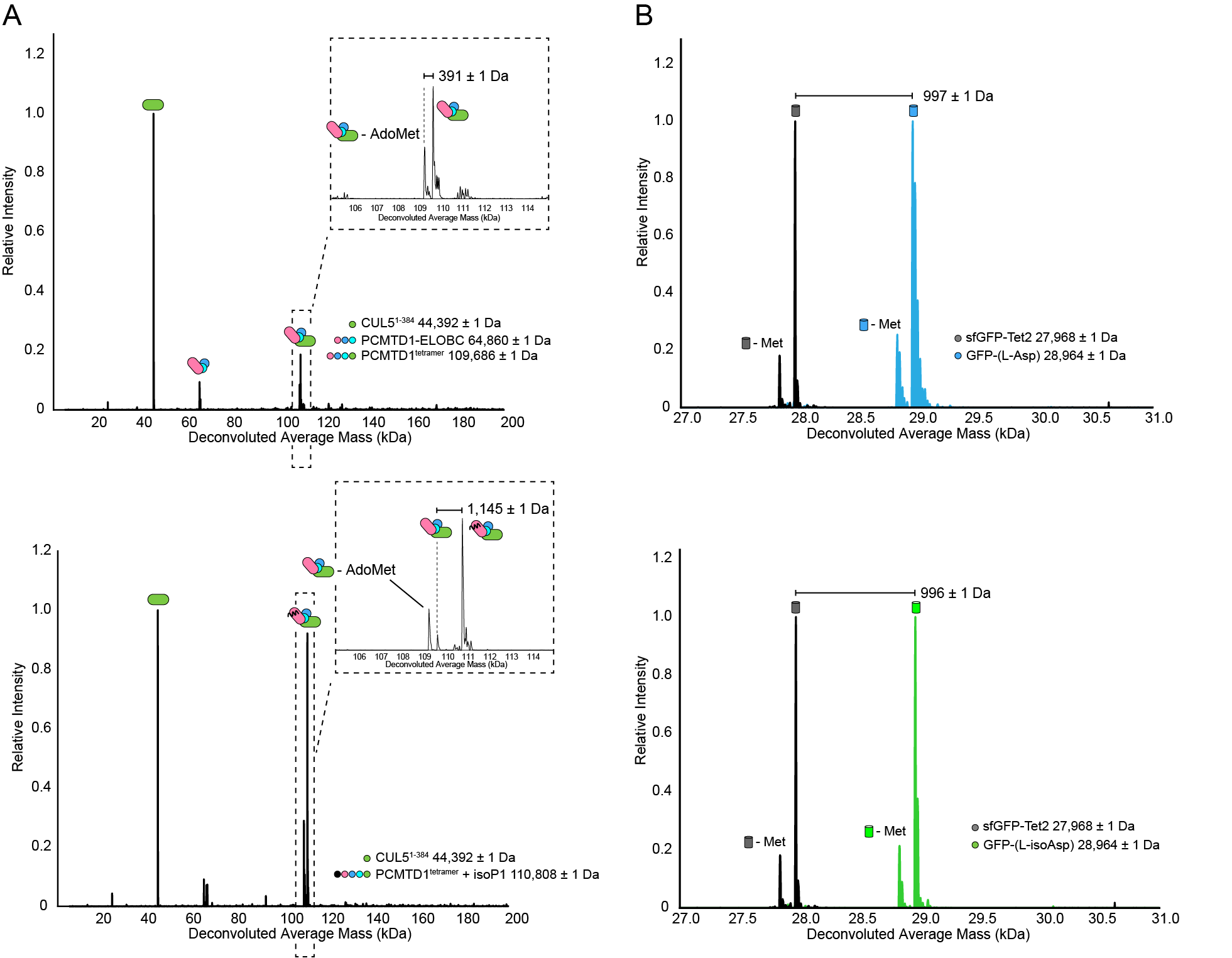

**Supplemental Figure 6. Native mass spectrometry ligand-binding controls, related to Figure 4.** A) Native mass spectra of PCMTD1-tetramer spiked with GGGKASADLAKY (top) and GGGKASA(L-isoD)LAKY (bottom). B) Native mass spectra of sfGFP-Tet2 before and after addition of TCO-modified peptides to generate sfGFP-Tet-TCO-GGGVYPDHA (GFP-L-Asp, top) and sfGFP-Tet-TCO-GGGVYP(L-isoD)HA (GFP-L-isoAsp, bottom).

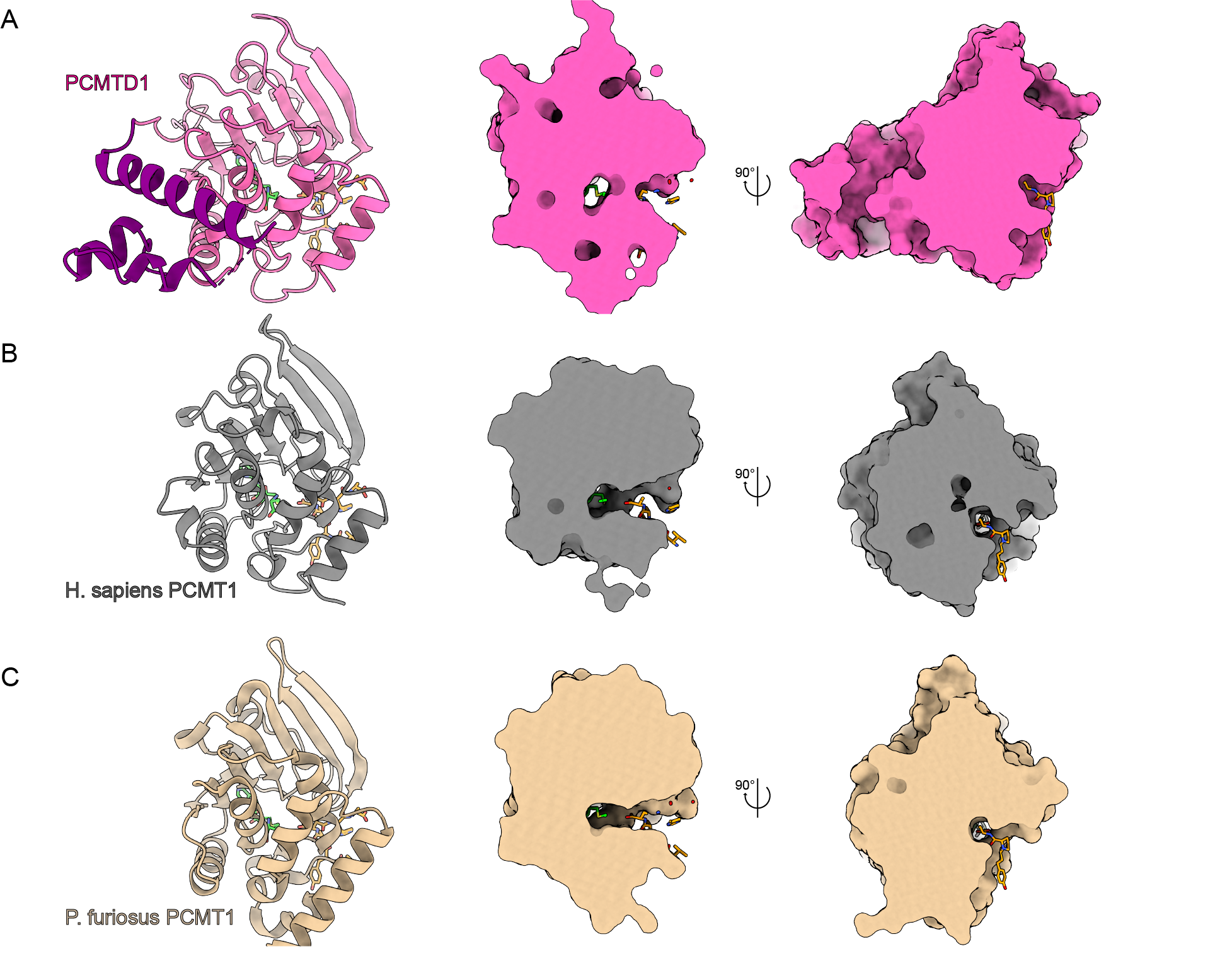

**Supplemental Figure 7. Cross sections of the co-factor and ligand binding sites of PCMTD1 and PCMT1, related to Figure 5.** Structures of (A) PCMTD1, (B) *H. sapiens* PCMT1, (C) and *P*. *furiosus* PCMT1 with surfaces shown. Positions for AdoHcy (lime; PDB IJG1) and VYP(L-isoD)HA (orange; PDB 1JG1) were generated by structural alignment of the crystal structure of *P. furiosus PCMT1* solved with ligands against the structures of PCMTD1 and *H. sapiens* PCMT1.

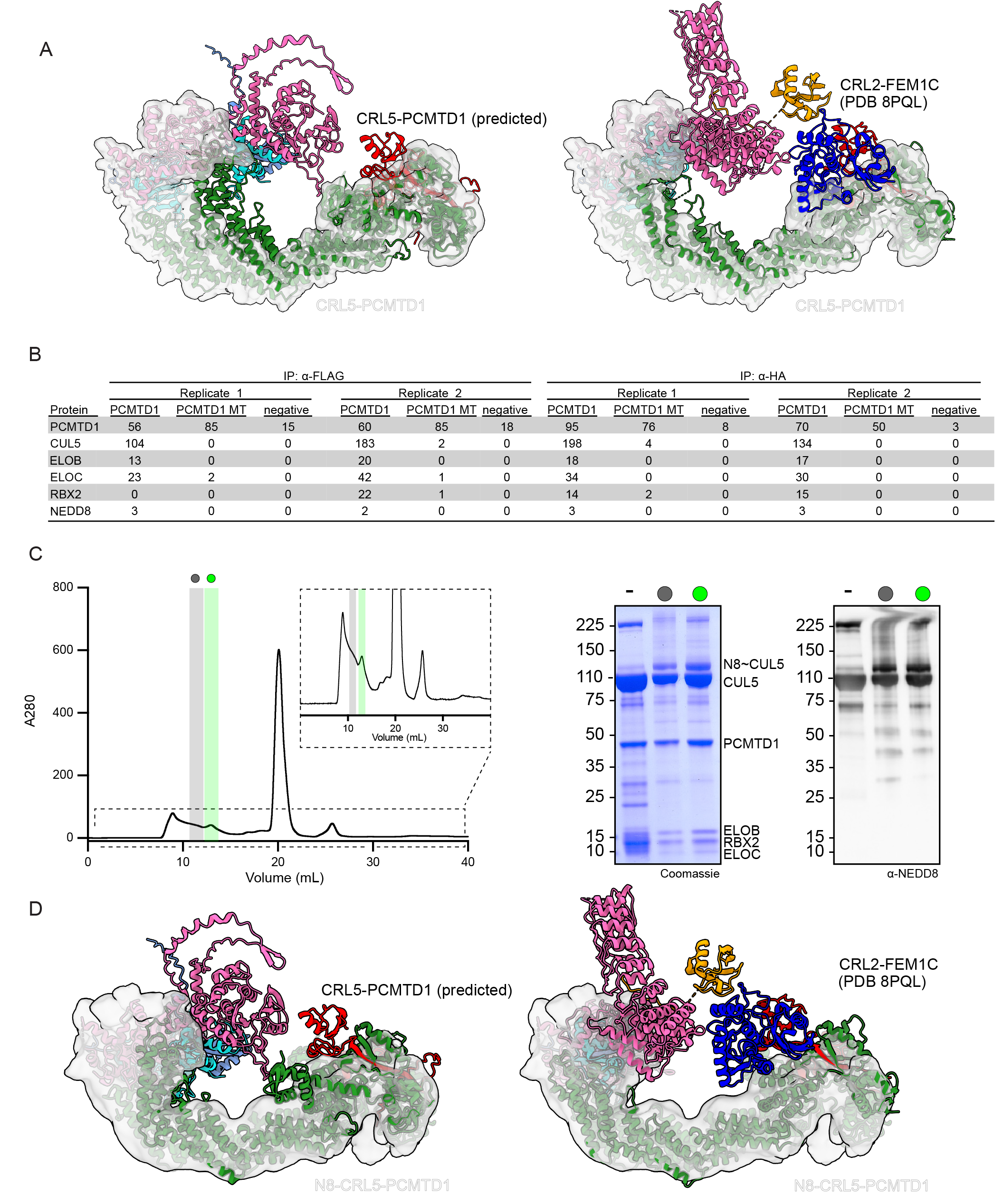

**Supplemental Figure 8. Sample preparation of N8-CRL5-PCMTD1, related to Figure 6.** A) Predicted structure of CRL5-PCMTD1 (left) and cryo-EM structure of active CRL2-FEM1C (right, PDB 8PQL) docked into the 3D reconstruction of CRL5-PCMTD1. B) Spectral counts of protein interacting partners to PCMTD1 from co-immunoprecipitation/LC-MS. PCMTD1 MT = PCMTD1 P243 F247; negative = FlpIn-293 lysate; α-HA = co-immunoprecipitation with HA beads; α-FLAG = co-immunoprecipitation with FLAG beads. C) Gel filtration enrichment of CRL5-PCMTD1 after *in vitro* neddylation (left) with corresponding α-NEDD8 immunoblot (right) of selected fractions. - = sample before neddylation. D) Predicted structure of CRL5-PCMTD1 (left) and cryo-EM structure of active CRL2-FEM1C (right) docked into the 3D reconstruction of neddylated CRL5-PCMTD1.

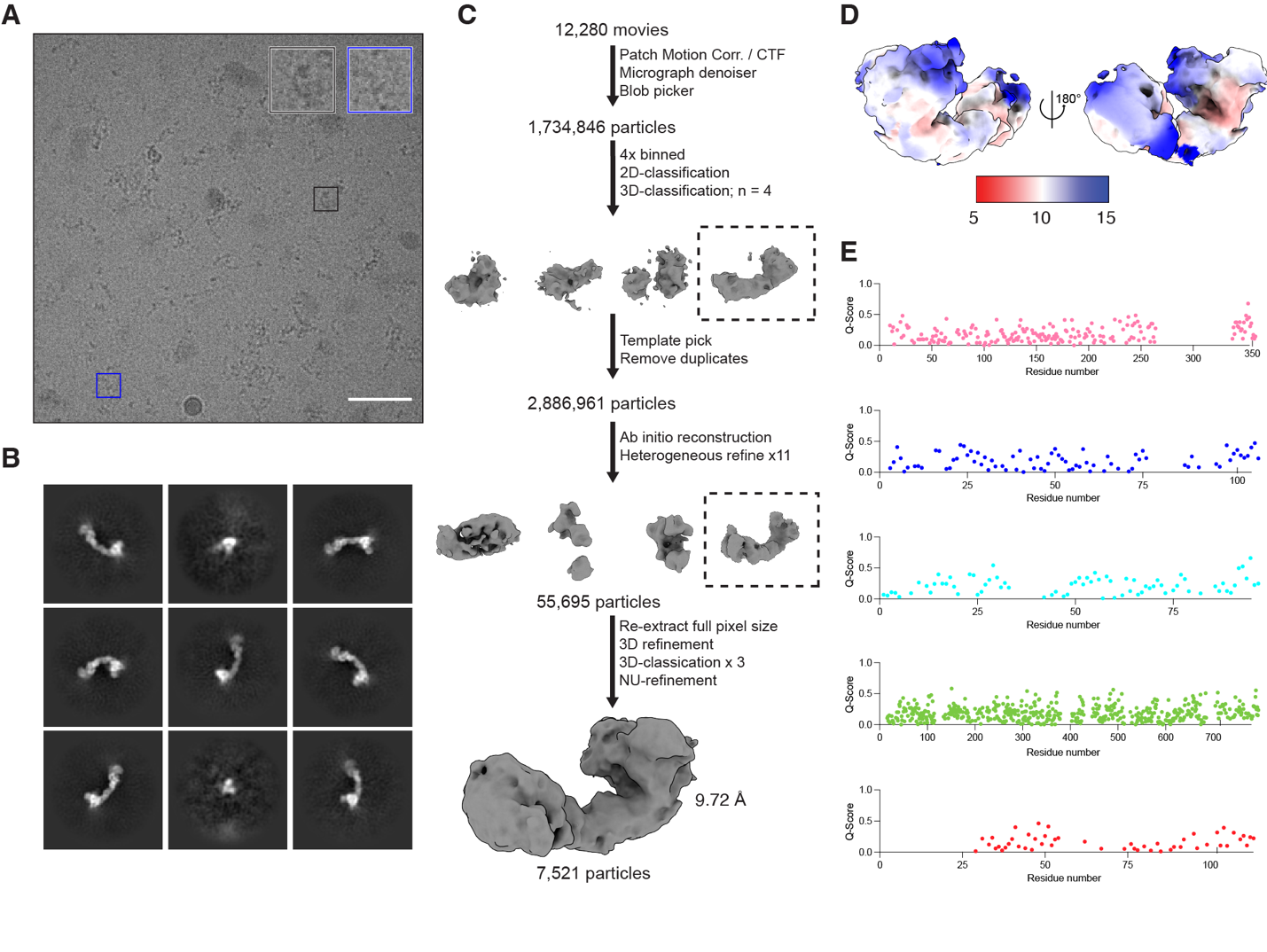

**Supplementary Figure 9. Cryo-EM 3D reconstruction pipeline for N8-CLR5-PCMTD1, related to Figure 6.** A) Cryo-EM micrographs of N8-CRL5-PCMTD1 after sample vitrification using the Chameleon system. Scale bar = 50 nm. B) 2D class averages of N8-CRL5-PCMTD1 from cryo-EM datasets. C) Data processing pipeline for CRL5-PCMTD1 with masks used for local refinement. D) Local resolution of the consensus map of N8-CRL5-PCMTD1. E) Q-score analysis of N8-CRL5-PCMTD1 model-to-map fit. Pink = PCMTD1; blue = ELOB; cyan = ELOC; green = CUL5; red = RBX2.
